# New insights into iodide metabolism based on preclinical models: impact on radiotherapy efficacy and protection against radioactive iodine exposure

**DOI:** 10.1101/2023.10.11.561684

**Authors:** Julien Guglielmi, Grégoire D’Andréa, Fanny Graslin, Kaouthar Chatti, Aurélie Schiazza, Sabine Lindenthal, Jacques Darcourt, Béatrice Cambien, Thierry Pourcher

## Abstract

**Background:** The main basic aspects of the regulation of thyroid metabolism by iodine are known, but given the complexity of the mechanisms involved, further analyzes in living animals are still required. Here, we provided new insights into iodine physiology but also into the optimization of radiotherapy with iodine, as well as effective countermeasures in the case of an exposure to radioactive iodine.

**Methods:** We performed Single Photon Emission Computed Tomography (SPECT) coupled to an X-ray scanner to record radiotracers in living mice and rats. Our imaging system was similar to that routinely used in nuclear medicine but was specifically designed for studies with small animals. Different modalities of administration of radioactive iodine or its radioactive analogues combined with a low or high iodine diet have been studied in pregnant, lactating and control animals. To optimize countermeasures against acute or chronic iodine exposure, the protective effects of potassium iodide (KI) administration protocols were analyzed. Perchlorate was administered to study the iodine metabolism in the kidney and stomach.

**Results:** Our results showed how the various organs capable of iodine uptake adapt to an iodine-deficient diet. Indeed, the uptake capacity of the thyroid gland, but also that of the salivary glands was significantly increased on a low iodine diet. In contrast, the iodine uptake capacity of the thyroid and lactating mammary glands was reduced on an iodide-rich diet. Our results also showed the physiological role of the kidneys in controlling excess circulating iodide. In addition, they revealed an active iodine cycle in the stomach. We also investigated the protective effects of daily KI administration during radioactive iodine exposure and found that the overall protection was better in rats (85%) than in mice (65%). We also included pregnant females and newborns, and we revealed the existence of specific mechanisms for the inhibition of the fetal thyroid by circulating iodine. Indeed, an iodine-rich diet or repeated daily administration of KI led to a strong inhibition of the iodide uptake capacity of the fetal thyroid.

**Conclusions:** Our study contributes to a better understanding of iodine metabolism and its regulation in the thyroid and in non-thyroidal organs in adult, fetal and newborn animals. Extrapolated to humans, our results not only provide better understanding of iodide withdrawal as a clinical preparatory measure for patients with differentiated thyroid cancer, but also help to optimize countermeasures in the case of an exposure to radioactive iodine.

## Introduction

Better knowledge of the regulation of iodine metabolism in the body is needed to optimize radioiodine therapy, and also to improve radiation protection during radioactive iodine exposure. The regulation of the iodine metabolism depends mainly on the level of circulating iodide. This in turn is determined by the dynamics of iodine uptake, accumulation and excretion in various organs including the thyroid, salivary glands, stomach, bladder and mammary glands. In females, pregnancy also has an effect on the level of circulating iodine.

Iodine is a component of the thyroid hormones 3,5,3 -Tri-iodo-L-thyronine (T3) and 3,5,3’,5’-Tetra-iodo-thyronine or thyroxine (T4). In the thyroid, these hormones are synthesized in several steps involving oxidation and protein iodination mechanisms confined to the colloid of the thyroid follicle. Iodine is a relatively rare element in the biosphere, but most mammals have the ability to efficiently absorb iodine from the blood into the thyroid. Therefore, the daily requirement for humans, for example, is about only one hundred micrograms.

Iodine metabolism in the thyroid is very finely regulated. This regulation is mainly based on active iodide transport mediated by the sodium/iodide symporter (NIS) ^1, 2^. The symporter is located in the basolateral membrane of thyrocytes where it mediates iodide uptake from the blood. Iodide then diffuses to the colloid where it is organified and incorporated into thyroglobulin, the precursor of thyroid hormones. The iodinated thyroglobulin is reabsorbed by the thyrocytes and converted into the hormones T4 and T3 which are then secreted into the bloodstream. During this process, part of the iodine is released and then recycled. At the physiological level, it has been clearly established that thyroid function is mainly regulated by Thyroid Stimulating Hormone (TSH) and iodine ^3^. This regulation of thyroid function involves, among other enzymatic activities, the capacity for NIS-mediated iodide uptake into thyrocytes. TSH stimulates the thyroid function through an enhanced ability to accumulate and organify iodide ^4^ and thus increase the secretion of thyroid hormones ^5^. TSH secretion by the pituitary gland is induced by a decrease in circulating T4-T3 levels (via TRH) ^6^. Interestingly, seve ral studies including ours ^7,8^ have demonstrated a direct feedback effect of elevated circulating thyroid hormone levels on the thyroid. Finally, the thyroid can adapt to highly variable iodine intakes from the diet. An increase in the concentration of circulating iodine, as can occur upon the ingestion of a food rich in iodine for example, will lead to a decrease in the thyroid avidity for iodide via a regulation of NIS activity ^4,9,10,11,12^. This regulation is also associated with a transient decrease in the secretion of thyroid hormones described as the Wolff-Chaikoff effect ^13^. So far, knowledge about the regulation of iodine metabolism by iodide is mainly limited to the cellular or organ levels. A study of this phenomenon at the scale of the whole organism is thus required for a better understanding of the dynamics between the various organs of capture and transit. Optimization of radioiodine therapy as well as protection against radioactive iodine exposures, may both directly benefit from this improved knowledge.

In current clinical practice, iodine-131 is used routinely as an adjuvant treatment for differentiated thyroid carcinoma (DTC). Treatment effectiveness is dependent on the amount of radioiodine that accumulates in the thyroid. It is now well established that iodine-131 radiotherapy is a very effective approach to treat most patients with differentiated or even metastatic cancer. However, a significant proportion of these patients develop refractory forms of cancer ^14^. In addition, adverse effects of RAI (radioiodine) therapy include chronic salivary dysfunction, which is observed in a substantial number of patients. Better understanding of the dynamics of radioiodine metabolism in the thyroid and salivary glands may help improve the benefit/risk balance.

During a nuclear power plant accident or atomic bomb explosion, large amounts of radioactive iodine can be released. The thyroid’s remarkable avidity for iodine, associated with its radiosensitivity, makes iodine a major element in radiotoxicology. The Chernobyl accident unfortunately demonstrated clearly, this with an increase in the frequency of thyroid cancers, especially in young exposed populations ^15,16,17^. However, administration of KI (potassium iodide) just prior to exposure is an effective protective measure against acute exposure to radioiodine. This KI intake is often possible in anticipation of the passage of the radioactive cloud emanating from a nuclear plant following an accident such as that of Chernobyl. KI intake (130 mg tablet in France) leads to a dilution of the radioactive isotopes, a saturation of the blood by iodide, and a decrease in the thyroid iodide transport capacity ^18^. It should also be kept in mind that the most radioactive iodine isotopes have relatively short half-lives (e.g., iodine-131, with a half-life of 8 days) and that long-term protection of a contaminated area is not the main concern in case of radioactive iodine contamination. In contrast, radioactive cesium, which is also released during a nuclear accident requires a more long-term protection, for example. Toxicology studies have indicated that a single intake of KI has not toxic effects and thus does not pose a health risk. Indeed, KI tablets were distributed in Poland during the Chernobyl accident and only a few very marginal cases of intolerance were noted ^19^. However, the Fukushima accident in 2011 has intensified the search for effective countermeasures against prolonged exposure. Indeed, the damaged plant released different isotopes over a period of about 10 days, resulting in repeated contaminations. A countermeasure that can be used in this case of chronic exposure and that is based on the daily intake of iodine has then been developed ^20^. To date, related studies essentially aimed at assessing dose-response of the protective effects on the thyroid ^21^, iodine pharmacokinetic parameters ^21,22^ as well as side effects ^23–27^. However, for optimal protection against repeated exposure, it is necessary to gain further insights into the dynamics of iodide capture and release between the various organs at the scale of the whole organism, and to investigate its regulation by KI. These studies should also include embryos and newborns. SPECT (Single Photon Emission Computed Tomography) imaging of small animals is a particularly suitable technology for these studies. It allows the dynamic and simultaneous monitoring of radiotracer accumulations in the different organs and thus the measurement of the potential protection provided by KI intake.

The aim of this preclinical study was to use our SPECT-CT imaging equipment to better characterize the iodine metabolism and its regulation by iodide in living adult, embryo and newborn rodents. In particular, we studied the influence of dietary iodide intake on radioiodine therapy and the daily KI administration as a countermeasure against accidental radioactive iodine exposure.

## Materials and methods

### Animals and SPECT imaging

The animals were C57BL/6J mice and Wistar rats. Iodine-124 or perchenetate-99mTc was administered intraperitoneally, orally, or by gavage according to the procedures described below. Radiotracer biodistribution images were obtained using a dedicated microSPECT/CT scanner (General Electric eXplore speCZT CT120 model) following procedures previously described ^7^. For each acquisition, the animal was placed in a gas anesthesia chamber (isoflurane, 2.5% for induction). Rodents were kept asleep by gas anesthesia (isoflurane, 1.5%) and placed in a thermostatically controlled bed for image acquisition. The bed was then positioned inside the scanner. A pulmonary ventilation control system was connected to a computer and used to monitor the animal’s respiratory system. Acquisitions were started at different times as indicated in the experiment procedures. The duration of the scans was set between 10 minutes and half an hour according to the signal intensity.

The reconstructed images were analyzed and quantified using AMIDE software. The absorptions were expressed as percentages of the injected activity after correction for radioactive decay. The radioactive decay correction allows the study of ratios to the initial administered iodide. Uncorrected signals allowed measurements of the remaining radioactivity at a given time relative to the initial dose.

The animals were treated in accordance with the French Agriculture Ministry guidelines, and the experiments were approved by the University Cote d’Azur Animal Care User and Ethics Committee (CIEPAL-Azur, references NCE/2014-211, APAFIS#19050-2019030815344840 (V3) and APAFIS#24082-2020021210254364 (V3)).

### Studies of the effects of animal pre-treatments on iodine accumulation capacity (Figures 1, 3, 5, 8, 9, and 10) using the biodistribution of pertechnetate-Tc99m

50 MBq pertechnetate-Tc99m was administered by intraperitoneal injection (Figures 1, 3, 5, 8, 9, and 10). SPECT image acquisitions were performed either one hour after radiotracer administration (Figures 1, 3, 5, 8, 9, and 10) or at different times after radiotracer administration (Figure 5, and Supplemental Figure 5). Then the animal was sacrificed by cervical dislocation. The increased accumulation capacity due to iodine deficiency (Figure 3) in mice and its inhibition by stable iodine administration in mice (Figures 8 and 9) and rats (Figure 10) were also studied. For the effects of pre-administration of KI on the regulation of iodine transport capacities by organs, measurements were performed after a 24-hour withdrawal from the last KI administration to minimize direct competitive effects. The effects of perchlorate administration aimed to study renal reabsorption of iodide.

**Figure 1:**
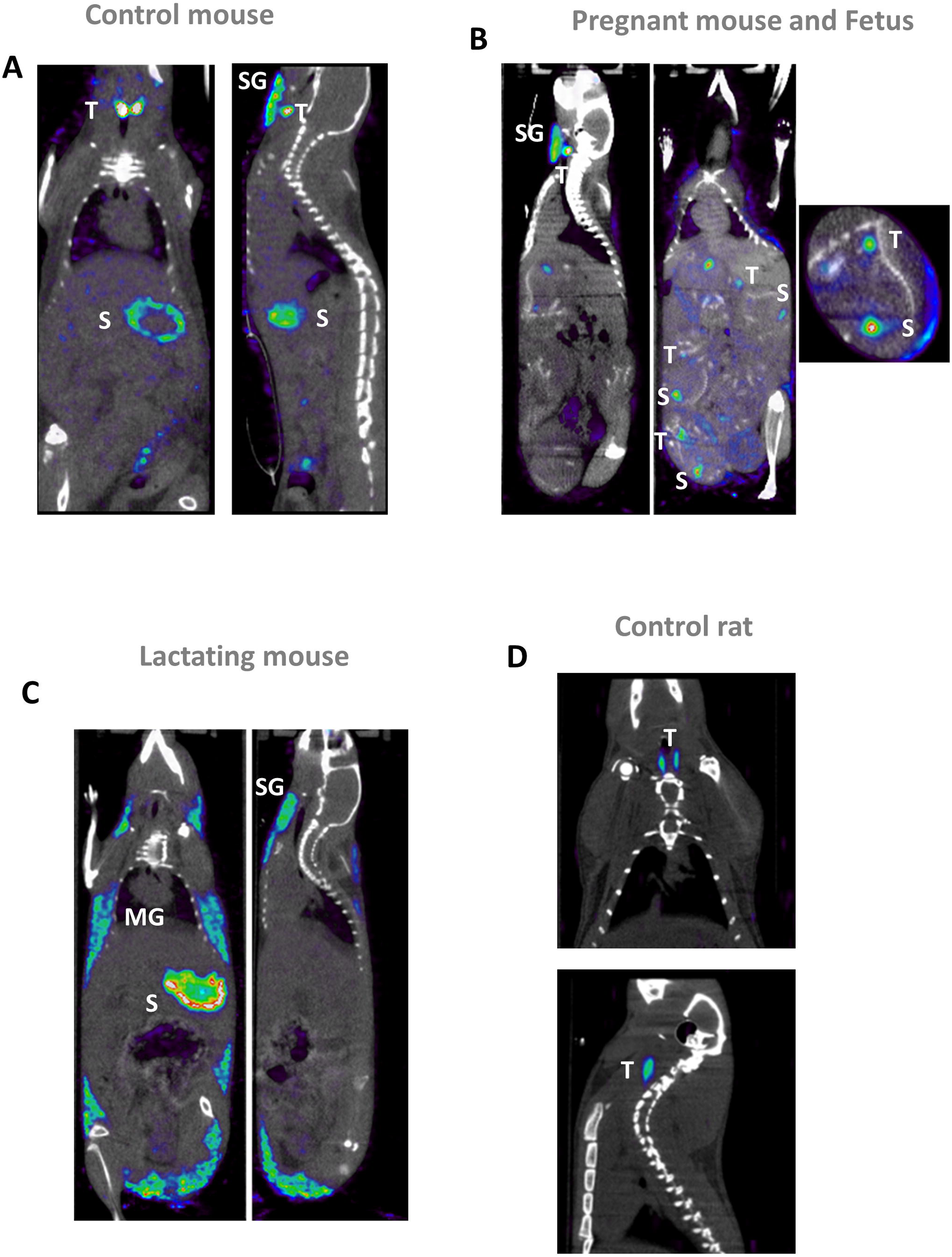
Representative SPECT-CT images of control (A), pregnant (B), lactating (C) mice or control rat (D) after intraperitoneal injection of pertechnetate-Tc99m. See study protocol in Supplementary Figure 1A. (T: thyroid; SG: salivary glands; S: stomach; MG: mammary glands).

**Figure 2:**
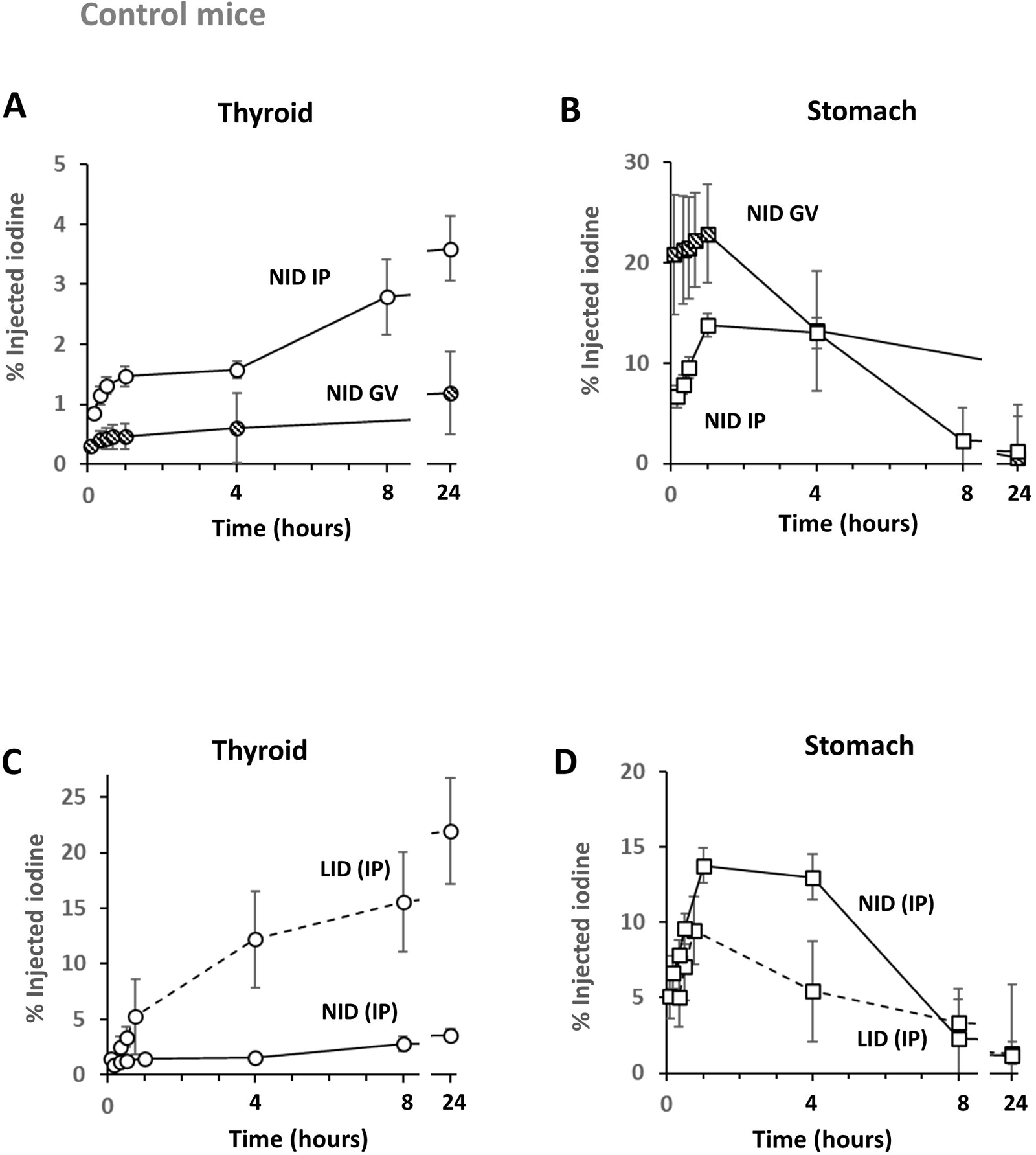
Kinetics of iodine-123 uptake in mouse organs: thyroid (A, C) and stomach (B, D), as a function of the mode of administration (A, B) and the iodine diet (C,D). Results obtained by intraperitoneal injection (IP) have white fill marks and those obtained by gavage (GV) have hatched marks. The kinetics plotted in dotted line are those obtained with mice under a Low­ iodine diet (LID) and in solid line those obtained with mice under a normal-iodine diet (NID). Values have been corrected for radioactive decay and are the average of the percentages of administered doses obtained from at least 5 mice. See study protocol in Supplementary Figure 1B.

**Figure 3:**
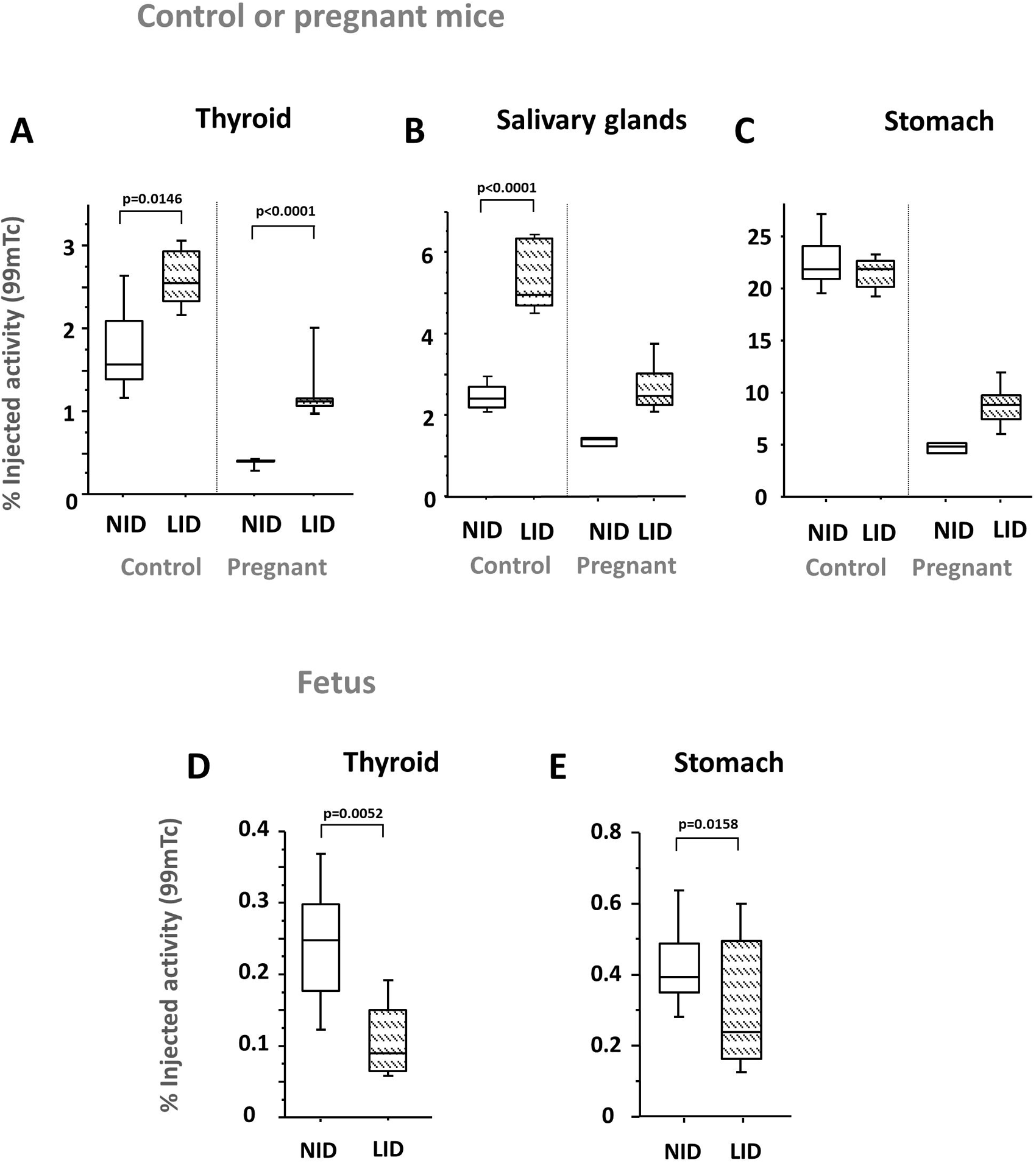
Effect of low-iodine diet (LID) on pertechnetate-Tc99m uptake in adult control and pregnant mice. Percentage of uptake of the injected dose obtained by organs on 5 adult control mice at 60 minutes after radiotracer injection. Values obtained with at least 5 mice. Measurements were made on the following organs of adult mice (control or pregnant): thyroid (A); stomach (B); salivary glands (C). In the embryo, only the thyroid and stomach were analyzed. See study protocol in Supplementary Figure lC.

**Figure 4:**
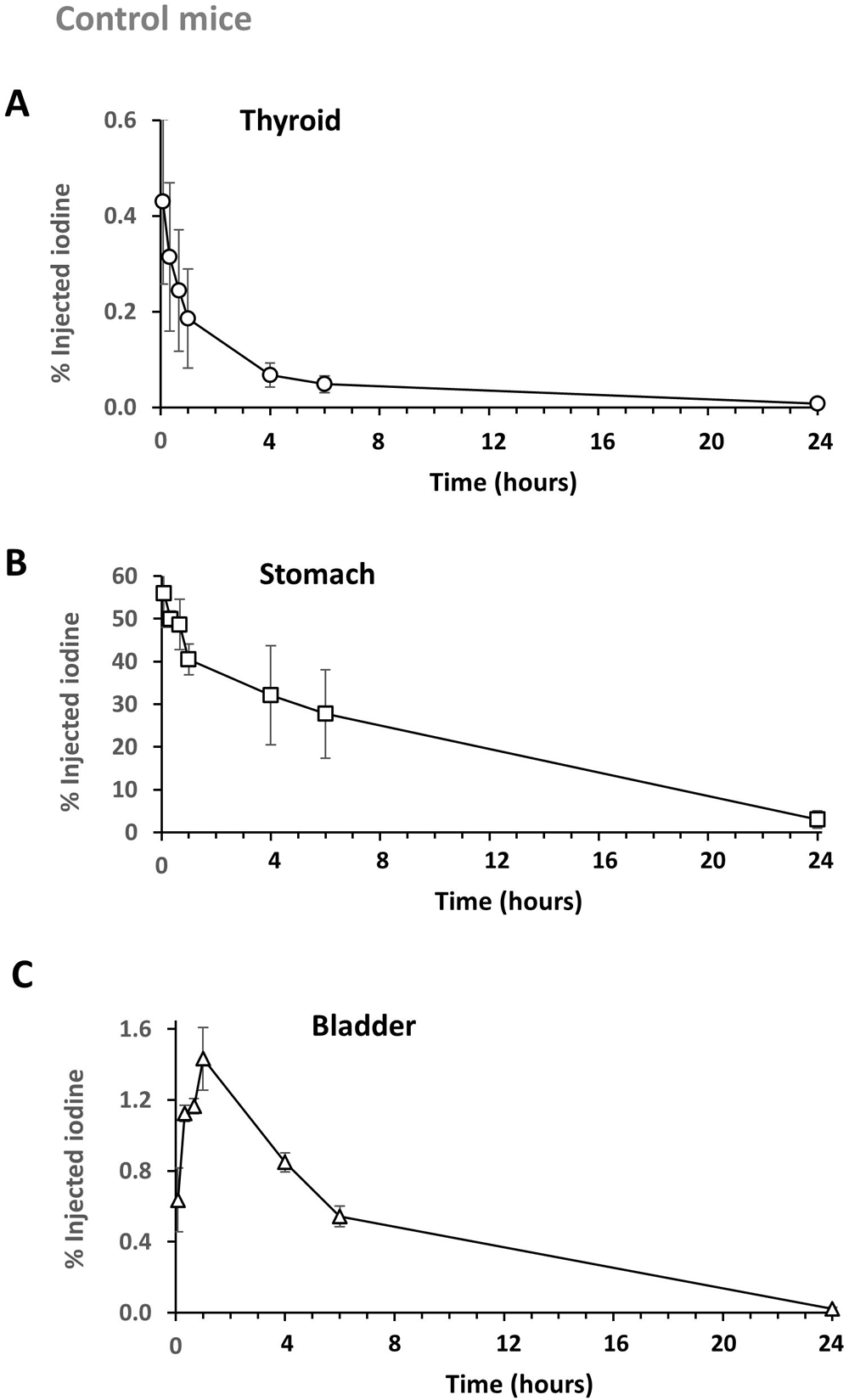
Kinetics in NID mice of iodide excess (1 mg/kg) administered by gavage. Kinetics were measured in the thyroid (A), stomach (B) and bladder (C). Values were obtained following co-administration of stable iodine and the radiotracer and are the averages of the percentages of injected doses obtained from at least 5 mice. See study protocol in Supplementary Figure 4A.

**Figure 5:**
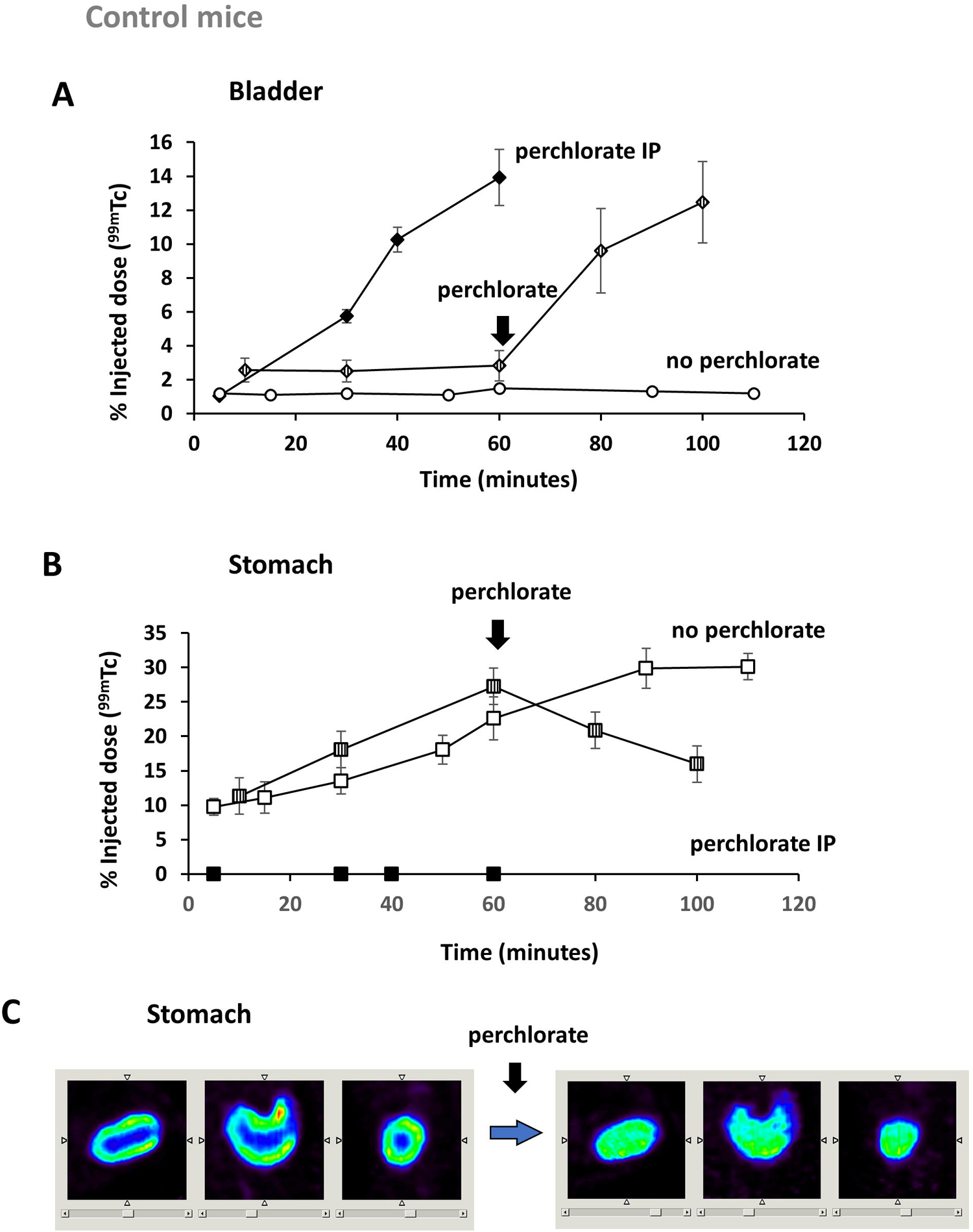
Effect of perchlorate administration on the kinetics of pertechnetate-Tc99m, administered by intraperitoneal injection. Values are the mean of the percentages of injected doses obtained in the bladder (A) and stomach (B) of mice. Values have been corrected for radioactive decay and are the means of the percentages of administered doses obtained from at least 5 mice. Representative SPECT images of pertechnetate-Tc99m accumulation in mice in the stomach (C). Animals were imaged 60 minutes after radiotracer injection (Tc99m) and immediately after perchlorate administration. See study protocol in Supplementary Figure 4B and 4C. Additional SPECT-CT images of the stomach area are available in Supplementary Figure 5.

**Figure 6:**
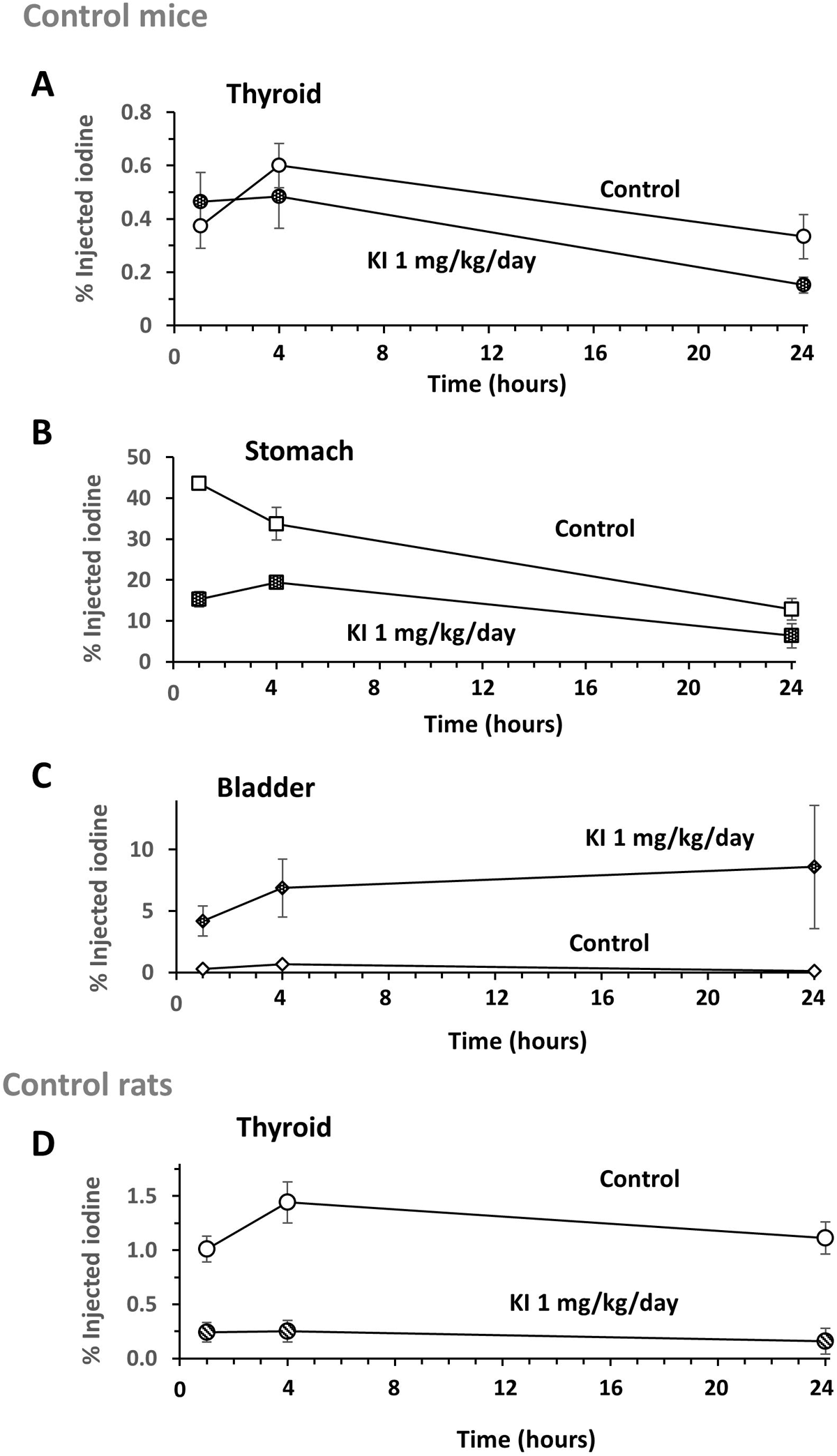
Effect in mice and rat of daily (over 5 days) gavage administration of iodine (lmg/kg) on the organ kinetics of iodine-123, administered by intraperitoneal injection. The values are the average from at least 5 animals of the percentages of the injected doses obtained in the thyroid (A), the stomach (B) and the bladder (C) of mice and the thyroid (D) of rats. They are compared to those obtained with control animals. See study protocol in Supplementary Figure 6.

**Figure 7:**
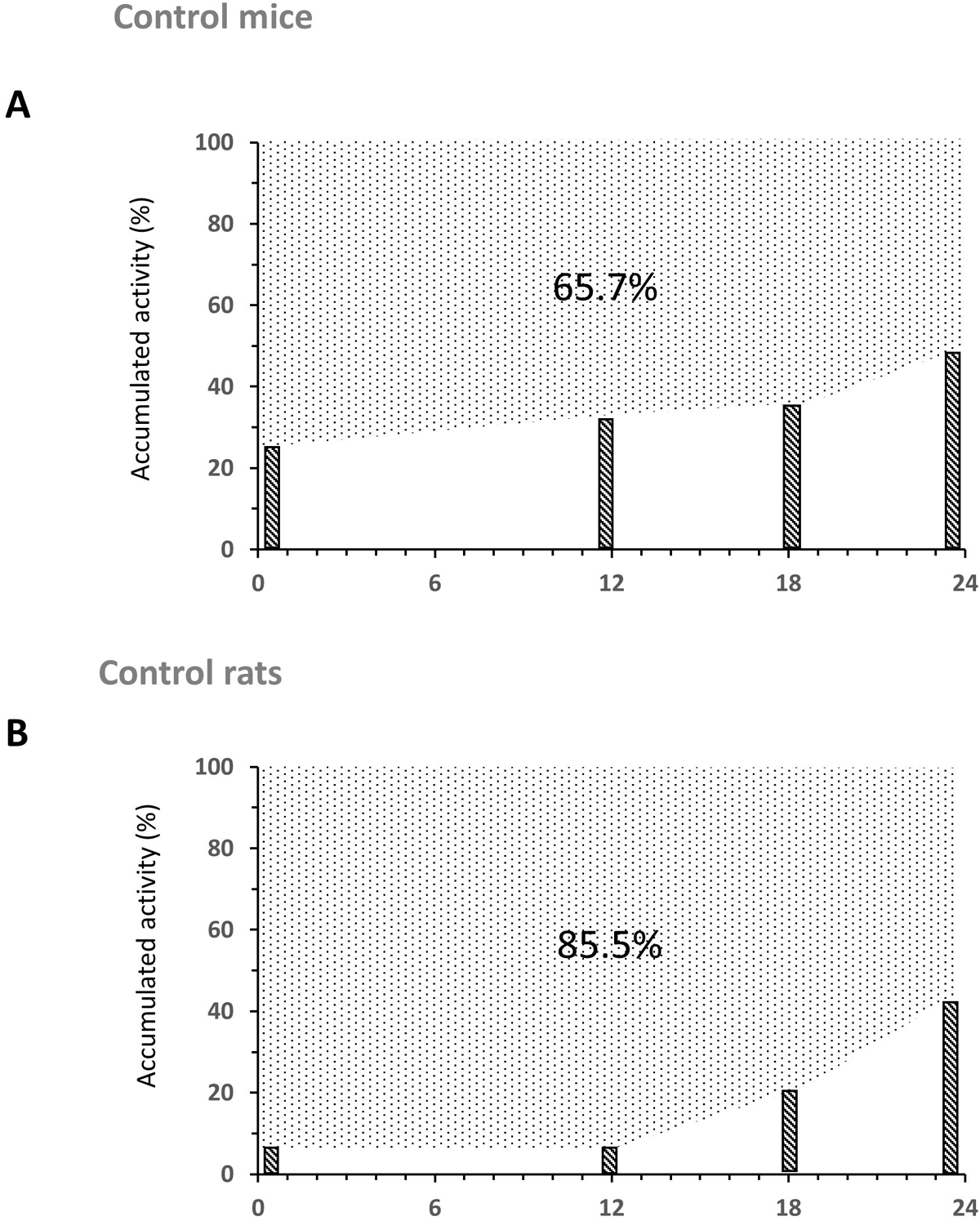
Estimation of protection of mice and rats thyroids by daily gavage (over 5 days) of iodine (1 mg/kg/day) following exposure to iodine 131. The dose was calculated from the administration of iodine 123 by intraperitoneal injection at different times between 2 intakes of Kl. Values are averages of thyroid dose calculations in mice (A) and rats (B) from at least 5 animals. See study protocol in Supplementary Figure 7A.

**Figure 8:**
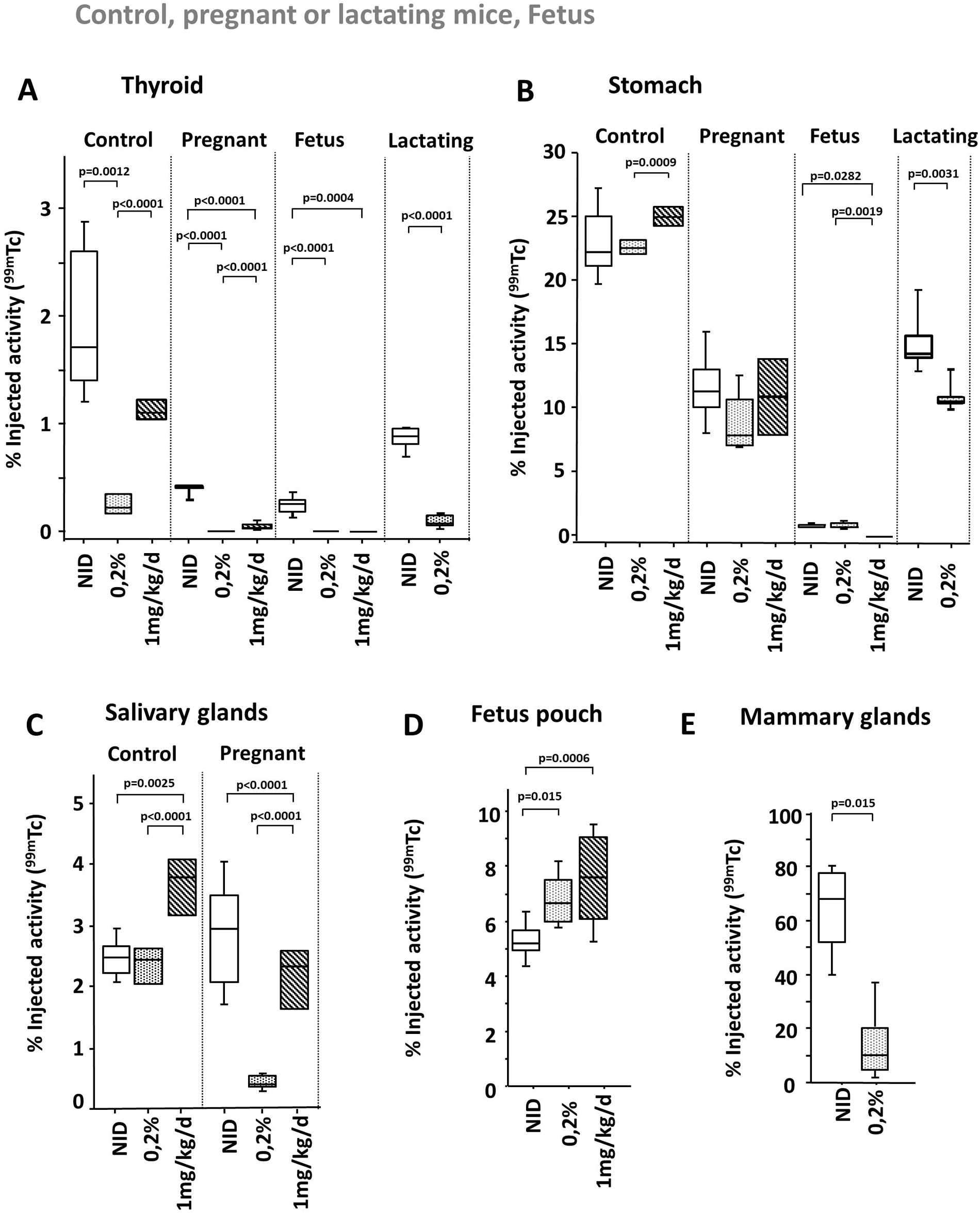
Effect of Rich-Iodide Diet (2 mg/Lin drinking water) or daily gavage (1 mg/kg/day over 5 days) on pertechnetate-Tc99m uptake in adult control, pregnant or lactating mice. A 24-hour weaning period was performed prior to measurement to limit a direct competitive inhibitory effect. Percentages obtained by organ (the thyroid (A), the stomach (B), the salivary glands (C), the embryo sac (D) and the mammary glands (E)) of uptake of the injected dose of at least 5 animals at 60 minutes after radiotracer administration. See study protocol in Supplementary Figure 8.

**Figure 9:**
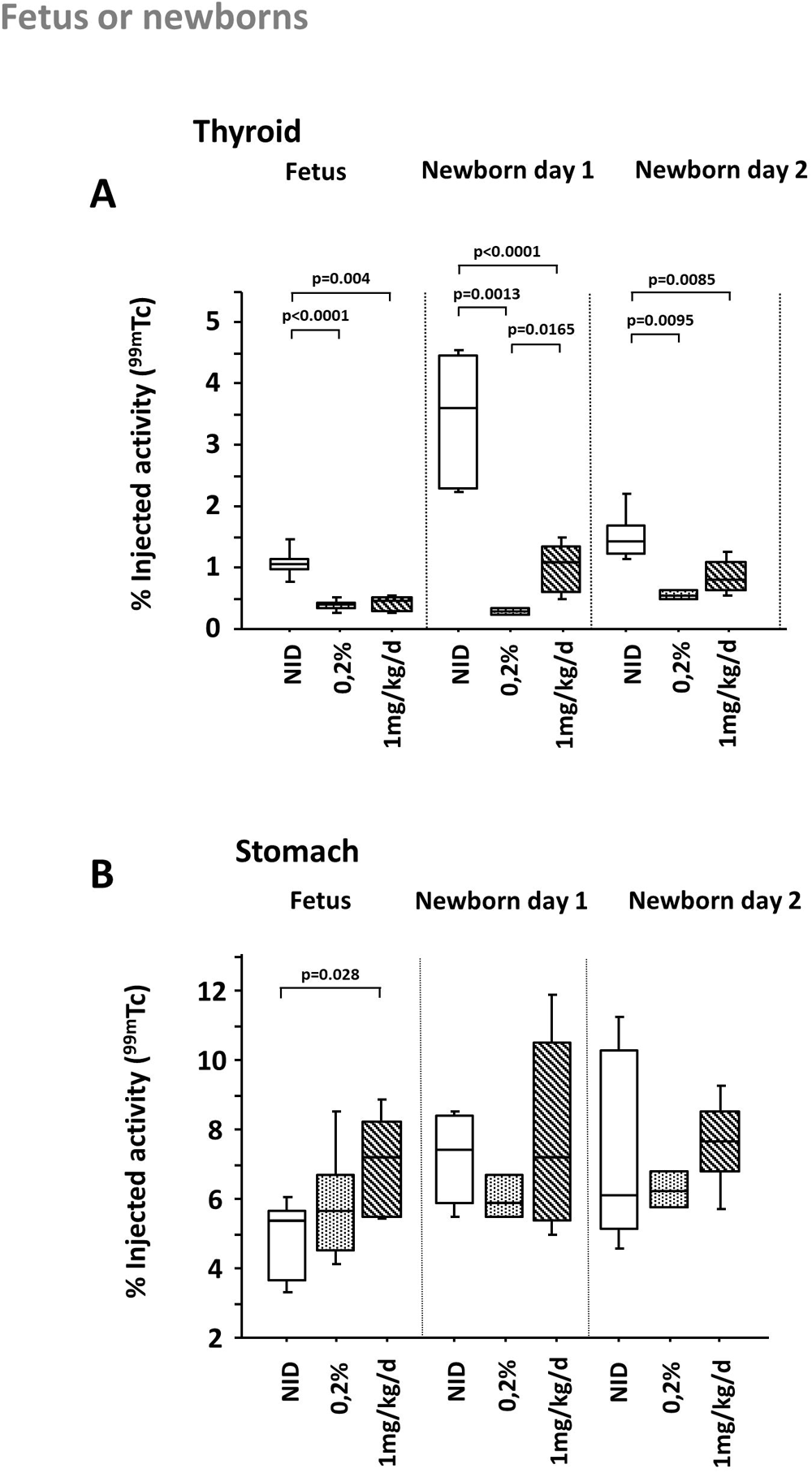
Effect of iodine-enriched diet (2 mg/L in drinking water) or daily gavage (1 mg/kg/day over 5 days) during maternal gestation on pertechnetate-Tc99m uptake in neonatal mice. A 24-hour weaning period was performed before the measurement. Percentages obtained by organ (thyroid (A) and stomach (B)) of uptake of the injected dose of at least 5 animals at 60 minutes after radiotracer administration. See study protocol in Supplementary Figure 8.

**Figure 10:**
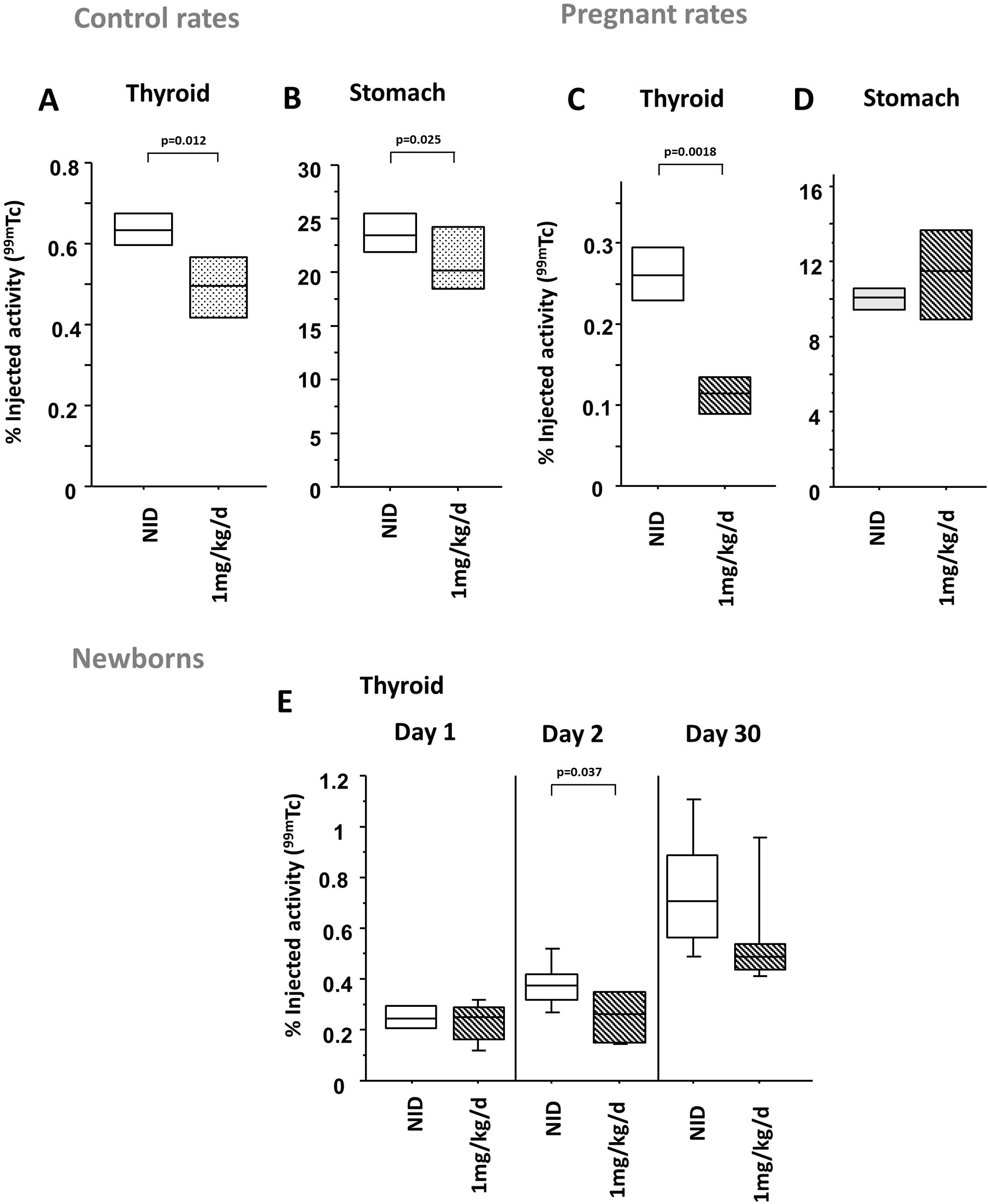
Effect of daily gavage (1 mg/kg/day over 5 days) during maternal gestation on pertechnetate-Tc99m uptake in control (A and B), pregnant (C and D) and neonatal (E) rats. A 24-hour weaning period was performed before measurement. Percentages obtained per organ (thyroid (A, C and E) and stomach (B and D)) of uptake of the injected dose at least 5 animals at 60 minutes after radiotracer administration. See study protocol in Supplementary Figure 8.

### Studies of the effects of pre-treatment of animals on iodine uptake and kinetics of acute contamination (Figures 2, 6, and 7) using biodistribution of Iodine-123

10 MBq iodine-123 was administered by intraperitoneal injection (Figures 2, 6, and 7) or gavage (Figure 2). SPECT image acquisitions were performed at different times after radiotracer administration. The scan durations were set between 10 minutes and half an hour depending on the signal intensity. After the first SPECT/CT imaging, the animal was placed back in its cage. For each SPECT imaging acquisition, the animal was anesthetized, imaged, awakened, and then returned to its cage. The procedure was repeated 24 hours after injection of iodine-123. Then the animal was sacrificed by cervical dislocation. These measurements allow the study of the iodine uptake capacity (accumulation and organification) by organ and thus to study modifications of these uptakes linked to a treatment. The experimental conditions are a priori close to the context of an acute contamination by radioactive iodine at the level of the molar concentrations of radioactive elements. The measurements are performed after a 24-hour withdrawal from the last KI administration in order to minimize direct competitive effects. We also studied the increased accumulation capacity due to iodine deficiency (Supplemental Figure 2) in mice and its inhibition by stable iodine administration (Figure 6) in mice and rats (Figure 7).

### Biodistribution of excess KI administration by gavage (Figure 4)

Animals were subjected to gavage administration of a cocktail of KI (1mg/kg) and iodine-123 (50 MBq) in a total volume of 200 µl. A series of SPECT imaging was performed at different times after gavage (1h, 4h, 6h, 24h). The purpose was to measure the biodistribution of excess iodine and its elimination kinetics.

### Biodistribution of pertechnetate-Tc99m as a tracer: Effects of perchlorate administration (competitive inhibitor) on biodistribution (Figure 5)

Animals were administered 50MBq pertechnetate-Tc99m by intraperitoneal injection (Figure 5). SPECT image acquisitions were performed at different times indicated after radiotracer administration. Perchlorate (2 mg per mouse) was administered as IP at the indicated times. Perchlorate administrations were aimed at studying renal reabsorption but also intra-stomach biodistribution.

### Simulation of protection by daily KI administration on the accumulated radioactivity dose according to different scenarios of accidental exposure to iodine-131 (Figure 7)

The animals were subjected to a repeated administration by gavage over 3 days of 1 mg/kg (weight of the mouse) of KI per 24h. We then studied the uptake (accumulation and incorporation) of 50 MBq of iodine-123 injected intraperitoneally at different times (0h, 12h, 18h and 23h) after the last intake of KI according to Supplementary Figure 7. We showed that this route ensures a calibrated dose and an almost instantaneous diffusion in the blood compartment (illustrated in Figure 2). This injection of iodine-123 was carried out on different animals at several times between the third and a possible fourth administration of KI: immediately after the third dose, 3h after, 6h, 12h, 18h or 23h after. A series of measurements on animals without KI administration was also performed. In all conditions, 1 hour after intraperitoneal injection of the radiotracer, we started the acquisition sequence by SPECT imaging. This procedure aimed to establish the effects of repeated KI intake on the decrease of the dose received by the rat or mouse thyroid in the face of radioactive iodine contamination. Doses for each administration of iodine-123 were derived from the integration of the areas under the curves of the amounts of radioactivity accumulated over 24h. Then, simulations of the radioactivity doses potentially delivered by iodine-131 were calculated from the iodine-123 kinetics. The percentage of protection was estimated from the difference between these results and the identical kinetics (Figure 7) obtained for control animals (without KI administration). The protection values (65.7% and 85.5% in Figure 7) was calculated as constant (compared to individual not protected by a KI excess) for all the times studied after the last intake of KI.

## Results

### SPECT imaging of adult, pregnant and lactating rodents

Figure 1A illustrates a representative SPECT-CT experiment in mice using pertecnetate-Tc99m showing the radiotracer accumulations in the thyroid, the salivary glands, and the stomach. These organs are the main areas accumulating iodide, but radiotracers can also be found concentrated in excretory organs such as the bladder. As illustrated in Figure 1B, SPECT-CT imaging of pregnant mice enabled the detection of the iodine uptake capacity in the thyroid and stomach of the embryos. In lactating mice, radiotracer accumulations were also found in the mammary glands (illustrated in Figure 1C). Experiments were also performed in rats and a representative result is illustrated in Figure 1D, where an accumulation of radiotracer was found in the thyroid and the stomach (not shown) but not in the salivary gland.

### Organ distribution and kinetics of iodine in mice according to different administration modalities

We measured over 24 hours the kinetics of iodide-123 accumulation in the organs of mice after intraperitoneal injection or after gavage of mice on a normal-iodine diet. Results for the thyroid and the stomach are shown in Figure 2A and 2B, respectively. The radioactive decay (half-life of iodine-123 is 13 hours) was integrated in the calculation of the results in order to measure the evolution of the amount of iodine administered in the different organs. In the thyroid (Figure 2A), a very rapid incorporation of radioactivity was observed with both modes of administration. This result indicates a very fast diffusion of the iodide-123 in the blood and then a fast capture by the thyroid. This rapid diffusion was expected with intraperitoneal injection but was also observed with gavage administration. For both modes of administration, thyroid uptake of the radiotracer increased with time up to 24 hours. Intraperitoneal administration resulted in 3-to 4-fold higher radiotracer levels than those obtained by gavage. We also studied the iodide-123 accumulation in the stomach with follow-up for 24 hours (Figures 2B). The administration of radiotracer by gavage to normo-iodinated mice resulted in rapidly decreasing kinetics after 8 hours with no detectable level after 24 hours. Intraperitoneal administration resulted in transient uptake in the stomach with a peak at just over one hour after injection and no detectable level after 24 hours. To a lesser extent, similar kinetics were observed in the salivary glands (Supplementary Figure 2C), with a similarly rapid and transient uptake.

### Effect of a low iodine diet on organ distribution and kinetics of iodine in mice

Iodine deficiency is known to increase the avidity of the thyroid for iodine. To study this phenomenon, animals were subjected to a normo-iodine diet (about 1µg of iodine per day) or a low iodine-diet (<0.01 µg of iodine per day) and imaged.

A 14-day iodine-deficient diet resulted in a significant increase in thyroid uptake with intraperitoneal injection of the radiotracer (Figure 2C). Regardless of the route of radiotracer administration, a large increase in uptake was also observed following iodine deprivation under gavage condition (see supplementary Figure 2A). In the stomach, only moderate decreases were induced by low-iodide diet (Figure 2D). Under gavage conditions (see supplementary Figure 2B), compared to mice on a normal-iodine diet, a higher retention was observed in the stomach of iodine-deficient mice which is probably linked to efficient mechanisms of organ retention (without organification). Altogether, these results suggested that there is no regulation of the iodide uptake in the stomach induced by a decrease in iodine concentrations and resulting lower levels in circulating thyroid hormones.

The iodine-123 SPECT experiments allowed simultaneous measurement of iodine accumulation and incorporation capacities, whereas the use of pertechnetate-Tc99m allowed measurement of NIS-mediated iodine uptake ^28, 29^. Tc99m has a higher energy than iodine-123 and enabled more reliable quantifications. However, its half-life time is shorter than iodine-123, which made measurements of long kinetics impossible. As shown by previous pertechnetate-Tc99m kinetics ^7^, changes in the organ capacities for active iodine transport, such as those induced by different iodine-level diets, can be determined by measurements 1 hour after radiotracer injection because a plateau is reached for the organs of interest. In order to achieve greater reproducibility of the administered doses, administration by intraperitoneal injection was also chosen for these measurements. As previously observed with iodine-123, the iodine-depleted diet resulted in an increase in iodine uptake capacity of the thyroids of both control and pregnant animals (Figure 3A). This increase in iodine uptake by the thyroid is expected to ensure thyroid hormone synthesis in case of iodide deficiency and to be induced by an increase in NIS expression ^30^. Figure 3C showed that, like in the thyroid, NIS activities in the salivary glands are increased under iodine-deficient conditions (see also iodine-123 kinetics in Supplementary Figure 2C) and associated with higher NIS protein expression levels measured by western-blot experiments (Supplementary Figure 3A). In contrast, no induction of iodine uptake in response to iodine deprivation was observed in the stomach (Figure 3B). This again indicates that NIS activity in this organ was not modulated by iodine.

Experiments in pregnant mice allowed us to measure iodine uptake in embryos. To our knowledge, we report SPECT experiment in mouse embryos for the first time. Figure 3D shows that the iodine-depleted diet resulted in a decreased iodine uptake capacity of the fetal thyroid, which contrasted with an increased uptake capacity of the maternal organ. This result indicates that induction by such a diet did not have the same effect on the embryo as on the mother and suggests that, in such iodine-deficient conditions, hormone production is likely to be reduced in the embryo. A decrease of iodine uptake capacity in the stomach was also observed (Figure 3E) and suggests, as for the thyroid, a lower transfer of the radiotracer from the mother to the fetus.

### Distribution and kinetics of acute excess iodide exposure by gavage in different organs of mice

Administration of KI pills is the basis of the preventive radiation protection strategy used for acute exposure. Repeated KI administration is the basis of the countermeasures against exposures over several days. Here, we aimed to establish the biodistribution kinetics of a large dose of iodide administered by gavage. For this purpose, acute excess KI (1 mg/kg per animal) was administered by gavage along with iodide-123. The results are shown for each organ in Figure 4. In the thyroid, the administered iodide-123 was quickly partially taken up but was also rapidly cleared as radioactivity was no longer detectable after 24 hours. This sharp decline is different from the slower kinetics observed when iodine-123 was administered alone as a radiotracer (Figure 2A). The kinetics were certainly related to fast saturation of the thyroid and the rapid elimination of the excess iodine. In the stomach, rapid decreases were also observed (Figure 4B). The excess iodide was excreted in urine and found in the bladder (Figure 4C).

### Effect of NIS competitive inhibitor on the pertechnetate elimination in urine and on stomach uptake

We observed that acute excess iodide exposure triggers elimination of iodide in urine, whereas iodide is not eliminated as quickly in the case of low circulating iodide concentrations encountered upon a normal-iodide diet or a low-iodide diet. This observation indicates an efficient iodide renal reabsorption for low circulating iodide concentrations. Therefore, we studied if the renal reabsorption of iodide could be mediated by the NIS-related protein. We used a NIS substrate, pertechnetate-Tc99m as a radiotracer, and a competitive inhibitor, perchlorate, whose efficacy on NIS activity is well established ^2, 31, 32^. Both radiotracer and inhibitor were injected intraperitoneally to induce a rapid passage into the blood compartment. As shown in Figure 5A, without the addition of perchlorate, the level of pertechnetate-Tc99m in the bladder remained very low (“no perchlorate” curve). In contrast, co-injection of perchlorate (“perchlorate IP”) immediately induced excretion of pertechnetate-Tc99m in the bladder by inhibition of its reabsorption. This inhibition of iodide reabsorption was also immediately observed upon perchlorate injection 60 minutes after administration of the radiotracer (Figure 5A, curve under the black arrow), indicating the existence of renal reabsorption by transporters with the functional specificities of NIS. Immunohistochemical analyses clearly identified an apical expression of NIS in some epithelial cells of the stomach (Supplemental Figure 5C). In the stomach (Figure 5B), the NIS-mediated accumulations were correlated with the circulating concentrations of pertechnetate-Tc99m that could be produced by the addition of a NIS inhibitor. Indeed, stomach pertechnetate-Tc99m accumulation was inhibited by perchlorate when injected early or after 1 hour. These inhibitions resulted in direct perchlorate-induced competitive inhibition of NIS-mediated gastric cell iodide accumulation together with elimination of the pertechnetate-Tc99m in the urine.

Figure 5C shows the biodistribution of pertechnetate-Tc99m in the stomach 60 minutes after intraperitoneal injection of the radiotracer and just after perchlorate administration. After 60 minutes (left SPECT images), the radiotracer mainly accumulated in the stomach wall. Low levels of radiotracer were found in the stomach lumen. The addition of perchlorate (Figure 5C) triggered an almost immediate distribution of the radiotracer in the gastric lumen, suggesting that the radiotracer uptake in the stomach wall is related to NIS-mediated reabsorption of iodide from the stomach cavity.

### Changes in the organ distribution and kinetics of iodine induced by previous daily excess iodine

In this set of experiments, we investigated the effects of daily excess iodide (1 mg/kg/day) on the organ distribution of iodide-123. Measurements were made after five daily administrations of KI followed by a 24-hour withdrawal period. According to our previous results (Figure 4), this period was sufficient to avoid a direct competitive effect of KI. We also assumed that this time was short enough to maintain transient effects, including inhibition of iodine accumulation capacity, for example. Daily KI administration resulted in a slight inhibition of iodine uptake in the thyroid (Figure 6A) and stomach (Figure 6B). As expected, iodine-123 was also detected in the bladder of mice after daily excess iodide but not in control mice (Figure 6C). This result indicates that daily KI administration increases the urine elimination of injected iodine-123.

Similar experiments were conducted in rats. Stronger inhibitions of iodine-123 thyroid uptake were observed in rats (Figure 6D) when compared to those found in mice (Figure 6A).

### Simulation of the overall protective effect of 1 mg/kg/day of KI against accidental exposure to radioactive iodine

Here, we aimed to simulate how a subject, who received a daily dose of 1 mg/kg of KI for four days, would be protected against acute radioactive exposure at different times after the last intake of KI. We therefore measured the accumulated radioactivity within thyroids after intraperitoneal administration of radioiodine at different times (0, 6, 12, 18 and 23 hours) after the last iodide intake. As iodine-131 is one of the main isotopes of risk in the nuclear industry, calculations were performed to simulate the accumulation of iodine-131 based on iodine-123 imaging. The results are presented in Figure 7 as the overall value calculated from the area under the maximum values for each time tested (0, 6, 12, 18, and 23 hours) compared with the value from a control experiment (without KI administration). The results obtained in mice indicate a protection of nearly 80% when the animals were exposed to radioactivity shortly after the last intake of KI. This protection decreased with increasing time between last KI intake and radiotracer administration, reaching a value of 50% by 24 hours. As shown in Figure 7A, the average area of the different protections obtained indicated an overall protection of 65%. Similar experiments were conducted in rats, with an initial protection of over 90% and an overall protection of 85% (Figure 7B).

### Effects of two iodide-rich diets, i.e., 2 mg/L KI in drinking water or daily gavage of 1 mg/kg of KI, on the organ uptakes of iodide in control, pregnant and lactating mice

The effects of excess iodide administration over five days on organ iodide uptakes were studied using SPECT imaging measurements with pertechnetate-Tc99m after a 24-hour withdraw to limit direct competitive effects. As illustrated in Figure 8A for control animals, iodine administration in drinking water induced an inhibition of uptake capacity in the thyroid. This inhibition was also found in pregnant (mother and fetus) and lactating animals. Such inhibition was more pronounced that in control animals that had been daily gavaged with 1mg/kg KI. For both iodide-rich diets, stronger and significant inhibitions of iodide uptake in the thyroid were observed in pregnant animals and especially in fetuses. Iodine uptake in the stomachs (Figure 8B) showed little effect of both iodide-rich diets. This observation indicated the absence of a direct competitive effect in our experimental conditions. The effects on the salivary glands (Figure 8C) were clearly unexpected, as the iodide uptake capacity was increased by daily gavage in control mice but decreased by iodide in the drinking water and by gavage (to a lesser extent) in pregnant females. In the fetus pouch (Figure 8D), moderate effects were induced by both iodide-rich diets. In lactating animals, only chronic administration in drinking water was studied and led to a strong uptake decrease in the mammary glands (Figure 8E). Western-blot experiments indicate that this decrease is linked to lower NIS protein expression (Supplementary Figure 3B).

### Effects of two iodide-rich diets, i.e., 2 mg/l KI in the drinking water or daily gavage of 1mg/kg KI, on the iodine uptake in organs of embryos and neonatal mice

Figure 9A shows the inhibitory effect of different iodine diets at the end of gestation on the 99mTc uptake capacity of the neonatal thyroid. In neonates, a transient hyperactivity was observed upon the first day after birth. This increase was strongly attenuated after treatment with chronic or daily excess iodine in drinking water or by gavage. No significant effect was observed in the stomach (Figure 9B), again suggesting that there is no inhibition by competitive action in our experimental conditions.

### Effect of the daily gavage of 1mg/kg of KI on the iodine uptake capacity in the control, pregnant and newborn rats

Administration of 1mg/kg per day in rats resulted in a mild but significant inhibition in thyroid uptakes similar to that observed in mice (Figure 10A). In contrast, only small differences in uptake capacity of the stomachs were observed (Figure 10B). This indicates that there is a moderate or no direct competitive effect by iodide. Administration of 1mg/kg per day induced a more pronounced inhibition of iodide transport capacity in the thyroid of pregnant rates (Figure 10C). In contrast to mice, few or no significant effects were observed on the thyroid of the pups (Figure 10E).

## Discussion

Optimization of radioiodine therapy as well as radiation protection against radioactive iodine exposures may both benefit from a better understanding of iodide metabolism and regulation within the body. Here, using dedicated SPECT-CT imaging to perform integrative studies in living animal models, we assessed the dynamics of iodine uptake/accumulation/elimination by the thyroid, salivary glands, stomach, bladder, and mammary glands in adult rodents as well as in embryos and newborns. We also examined how iodine metabolism is affected by different iodide-level diets and repeated administrations of KI, both measures having consequences in the clinical and radiation protection contexts.

The use of 3D tomographic SPECT-CT imaging permits the measurement of iodine uptakes in major organs of living adult mice, and to distinguish the uptake of organs in close proximity such as mouse thyroids and salivary glands ^7,29,33^. Combining two radiotracers allows to assess both the radioiodine uptake capacity (with pertechnetate-Tc99m) and its organification (ability to incorporate radioiodine in hormone synthesis using iodine-123). Thanks to a millimeter resolution, this technology has allowed us to perform intra-organ analyses and to visualize heterogeneous Na/I symporter (NIS) expression within mouse xenografts ^34^. Using a pin-hole collimator enabled us to perform, for the first time to our knowledge, uptake measurements in mouse embryos. Such detection is however not feasible in rats, for which a slit collimator only allows poor resolution.

### Information on thyroid physiology from rodent models

Rat models are often used to study thyroid physiology based on their ability to uptake radioiodine. However, it is well established that the salivary glands do not mediate iodide uptake in rats unlike in mice or humans. In addition, in the rat thyroids, our results showed that daily iodide administration evokes mild inhibition of pertechnetate-Tc99m uptake capacity (Figure 10A) but strong inhibition (Figure 6D) of iodide-123 (uptake and organification) and efficient protection (Figure 7B). The excess iodine-evoked inhibition of the organification in the rat thyroid has been already reported ^13,35–37^. In the mouse thyroids, daily iodide administration evokes similar mild inhibition of pertechnetate-Tc99m uptake capacity (Figure 8A) but mild inhibition of iodide-123 (Figure 6A) and less efficient protection (Figure 7A). These results indicate that an inhibition of the ability to organify iodine into thyroid hormones occurred in the rat thyroid but to a much lesser extent (or not at all) in mice. To our knowledge, only few reports suggested that iodide administration blocks iodine organification in the human thyroid of normal adult ^38,39^. From the thyroid point of view, rats could represent a more representative model of the iodide-evoked regulation of the human thyroid function compared to mice.

In contrast, mice and humans share similarities in NIS expression patterns and in iodide uptake ability in both the thyroids and the salivary glands, which suggest that mouse models may better recapitulate the global iodine metabolism in humans.

These differences in animal models must be considered when studying the iodide metabolism, the dynamics of radioiodine metabolism within the body as well as the inhibition of radioiodine incorporation by protection countermeasures.

### Repercussions of our thyroid findings on radiotherapy

The influence of the route of radiotracer administration on radioiodine efficiency has been explored by our group and discussed in the context of treatment of thyroid metastases with radiotherapy ^7^. Whatever the administration modality (oral and intraperitoneal injection), a similar behavior to iodine-123 was observed. The initial diffusion of the radiotracer in the thyroid via the blood compartment is very rapid. However, the iodide uptake in thyroid cells is 2-3 times lower for oral administration (see Figure 2A). In clinical practice, to protect the caregiver from radiation, iodine-131 is administered to the patient orally but injected doses should increase the dose. While it is well established that oral iodine-131 radiotherapy is a very effective approach to treat most patients with differentiated cancer (expressing NIS and mediating iodide uptake) and even patients presenting metastases. However, a significant proportion of these patients with differentiated cancer develop refractory forms of cancer ^14^. Early detection of refractory forms could allow to perform injection of iodine-131 aiming higher initial dose of iodine-131, as part of a precision medicine strategy.

Beside the route of radioiodine administration, a low-iodine diet was shown to drastically influence thyroid uptake in our studies (see Supplementary Figure 2). This increase in thyroid activity is likely to compensate for a decrease in iodine intake. Mechanistically, iodine deprivation has already been shown to increase NIS expression within thyroids in control rats or mice ^40^ ^7^, pregnant rats ^30^ and in humans during clinical studies ^41^. An increasing number of nuclear medicine departments are advocating a strict iodine-depleted diet prior to radiotherapy treatment, particularly for cases with a potentially high risk of recurrence.

An additional strategy against refractory differentiated thyroid cancer aims to improve NIS expression and NIS-mediated iodide uptake. In this context, the understanding of the molecular mechanisms leading to a decrease in NIS expression in refractory cancers ^42^ and the control of associated signaling pathways constitute a great hope for these patients ^43, 44, 45^.

### Integration of thyroid data for radiation protection

After the Chernobyl nuclear accident, epidemiological studies revealed that the main related risk was thyroid cancer in children living near the nuclear plant ^15, 46^. Referring to the increase of the NIS-mediated iodide uptake evoked by low-iodine diet (Supplementary Figure 2), the absorbed thyroid dose of people living in an iodine-deficient region is therefore a priori increased. Iodine deficiency in the populations of the Chernobyl region was certainly an aggravating factor in the rate of thyroid cancers seen in exposed children ^47,48^. For acute exposure to radioactive iodide, stable iodide prophylaxis (Iodine Thyroid Blocking) appeared as an easy and efficient method to limit radioactive accumulation in the thyroid and was recommended by the World Health Organization ^49, 50^. However, the Fukushima Daiichi nuclear accident differed in that there were several radioactive releases over 4 weeks, which introduced new challenges in radioprotection. As no recommendation exists in that context, our group sought to determine how daily administration of stable iodide could protect a population chronically exposed to radioactive iodine ^20^.

In our experiments, the mouse thyroids still contained detectable radiotracer 24 hours after radioiodine exposure. This retention is due to the unique ability of thyroids to incorporate iodine-123 in hormone synthesis. In contrast, radioactivity is quickly washout in the stomach and salivary glands (Figure 2 and Supplementary Figure 2). These results corroborate strategies focusing on specific thyroid protection over time in the context of accidental exposure to radioactive iodine.

To study countermeasures against radioiodine contaminations, percentages of the administered activities per organ should be established as an individual dosimetry for each organ. Here, oral administration of the radioiodine allowed to study its biodistribution in the context of a contamination by food. As for an intraperitoneal injection, fast absorption was observed after oral administration and radioiodine was rapidly taken up by the thyroid. Whatever the contamination route, these data suggest a rapid passage via the blood compartment into the organs. Intraperitoneal injections were therefore chosen to administer accurate quantities of radiotracer to rodents.

Then, we first evaluated the efficiency of long-term protection (case of repeated exposures) based on daily 1 mg/kg KI administrations. This procedure and the doses were chosen according to studies on plasma and urine iodine concentrations after a single KI administration ^21^ or repeated KI administration ^22^. Here, our results using SPECT-CT on the kinetics of excess iodide (1 mg/kg) administered by gavage (Figure 4) are correlated to those obtained by inductively coupled plasma mass spectrometry ^21^.

As expected from the kinetics of excess iodide indicating a strong reduction of circulating iodide within 12 hours, the inhibition of thyroid radioactive uptake fluctuates over time following daily KI administration. Consistent with a direct competitive effect of a circulating iodide excess, lower thyroid uptakes were measured right after the KI administration and higher levels prior to the next KI treatment. Initial protections measured within 30 minutes after KI administration reached up to 75% in mice and 90% in rats, respectively. By integrating thyroid iodine-123 content over 24 hours following five-day KI treatment, a global protection of 65% and 85% were estimated in mice and rats, respectively (Figure 7). The observation that the adaptive response to chronic iodine overload is more pronounced in rats than in mice is consistent with the aforementioned difference between rat and mouse in the organification regulation. The higher protection in rats is related to stronger iodine-evoked inhibition of organification. Such differences in thyroid regulations between both rodent models may have repercussions for biodistribution kinetics and protection, and thus corroborate the idea that several animal models may need to be used to get relevant results.

Alternatively to daily oral KI treatments, chronic KI administration in the drinking water (0.2 % of KI) was tested, following Eng et al ^11^. After five days of treatments, strong inhibitions were observed in the thyroids of control, pregnant and lactating mice but also fetuses. Such uptake inhibitions were higher than with oral KI treatments (Figure 8). Considering the associated adverse effects of such thyroid blockade, our results tend to designate oral KI administration as the most appropriate countermeasure for multiple putative exposures to radioactive iodide, offering a good compromise between protection and adverse effects. Consistent with our observations, tests in the literature on different species show an absence of undesirable effects of daily KI administration in adult or young models ^23,24^.

### Integration of salivary gland data for protection in the context of radiotherapy

No increase in salivary gland disease was reported in the epidemiological studies following the Chernobyl incidents. During radioiodine therapy, however, ^131^I accumulates in cells of salivary glands and compromises their function ^51^. The risk of developing secondary cancer, while low, remains debated ^52–55^. Xerostomia is, however, a frequent and often persistent complaint of patients. Despite the adoption of standard preventive measures, such as increased water intake, pharmaceutical methods, or repeated massages of the salivary glands ^56,57^, parenchymal damage and chronic salivary dysfunction are observed in a substantial number of patients. Better understanding of the dynamics of radioiodine metabolism in salivary glands may help reduce its uptake, increase its secretion and offer protection.

Our data (illustrated in Figure 1) showed that iodide uptake by the salivary glands is observed in mouse models, but not in rats, as expected from the literature ^58^. In humans, iodine accumulations in the salivary glands are mainly observed in the parotid and submandibular glands ^59^. Expression of NIS has been shown in salivary glands at the basolateral membrane of ductal cells ^60,61^. As previously described ^7^ and recommended by the American Thyroid Association ^56^, iodine deprivation is an important factor to improve the efficacy of iodine-131 radiotherapy by removing healthy thyroid residues and thyroid metastases. As in the thyroid, our results indicated that a low-iodide diet induced an increase in the iodine accumulation capacity of the salivary glands (Figure 3). This should be taken into account during radiotherapy of metastases when low-iodide diet is performed to improve therapeutic efficiency. This diet could be especially used when a refractory form is suspected. Then, for radioprotection reasons, such strict iodine-deficient diet should be combined with an induction of salivary secretion during radiotherapy to avoid side effects on the salivary glands.

### Insights gained into iodine metabolism from studies on the stomach

As for salivary glands, studying the effects of radioactive exposure of the stomach in the context of radiation protection is not relevant considering that no major incidence of gastric cancer was reported in the epidemiological studies following the Chernobyl incident. Stomach exposure is not an issue during radioiodine therapy either. Nonetheless, information on the role of the stomach in the management of iodine metabolism is still rather scarce. Our SPECT studies provide the opportunity to better understand the dynamics of iodine within gastric compartments and in the context of a whole organism.

In the stomach, iodine-123 delivered through intraperitoneal injection is rapidly accumulated from the blood into the stomach. This is consistent with increased uptakes of pertechnetate-Tc99m previously described ^7^. As evidenced by the submillimeter-resolution images obtained by SPECT /CT, the radiotracer uptake is mainly located in the stomach wall. This accumulation is catalyzed by gastric cells that express NIS in their basolateral membrane ^62,63^. Despite an expected diffusion of radiotracers towards the stomach lumen, radiotracers were still mainly observed in the stomach wall 60 minutes after injection suggesting either very weak diffusion or reabsorption from the lumen (Figures 1 and 5, Supplementary Figure 5). To choose between these two hypotheses, we administered perchlorate, a competitive inhibitor of NIS-mediated iodine transport. Perchlorate induced an almost immediate change in the localization of the radiotracer from the wall to the gastric lumen highlighting a mechanism of iodine reabsorption from the stomach lumen (Figure 5C). Then, we performed immunohistochemical analyses aiming to show if NIS located in the apical membrane of stomach cells could mediate such iodide transport. Our results clearly identified an apical expression of NIS in epithelial cells of the pylorus aera of the stomach (Supplementary Figure 5C). Taken together, our data show the existence of an active iodide cycling mechanism in the stomach. Circulating iodide is taken up and concentrated in the cells of the gastric and pyloric gland via the NIS protein located in their basolateral membranes. From the apical membrane of these same cells, iodide diffuses in part into the gastric cavity. Then, iodide is efficiently reabsorbed by other cells of the pylorus aera by active mechanisms involving the NIS protein located in the apical membrane of these cells. The iodide then returns to the blood compartment. This active cycling of iodide could be related to the antioxidant properties of iodine in its ionic form (I-) which would accumulate to protect cells against oxidative stress ^64, 65^. Conversely, reactive forms of iodine (e.g., I2) or other oxidized anions carried by NIS, such as thiocyanate ^66^ and nitrate ^67^, could have a bactericidal effect on the food bolus. One should note that NIS can not mediate reabsorption of oxidized forms of iodine as well as iodide, and then should be accumulated in the lumen. In any case, for the first time to our knowledge, this study highlights the dynamic cycling of iodine in the stomach and indicates a physiological role of iodine in the stomach, which still needs to be further characterized.

### Repercussions of renal iodine reabsorption in the context of radiation protection

Among patients treated for thyroid cancers, radioiodine therapy has not been associated with increased risk of developing secondary renal cancer. Getting better insight into renal iodine elimination, however, is key to improve the radioprotection of other organs. Therefore, urine elimination of radioiodine was studied by monitoring the radioactive content in the bladder after KI or perchlorate administration.

Without KI treatment, radioiodine is reabsorbed by kidneys and the bladder radiotracer content is low (Figure 6C). A single KI administration induced a rapid increase in the bladder radioiodine content reflecting the fast elimination of excess radioiodine from all extra-thyroidal organs following saturation of renal reabsorption mechanisms. A return to a non-saturating concentration of KI between 12 and 16 hours after KI administration is observed (Figure 4C). Administration at intervals of less than 12 hours is expected to result in continuous saturation of renal reabsorption and high level of circulating iodide with a potentially increased risk of toxicity. These measurements are in agreement with ICP-MS results in rats ^21,22^. Here, experiments with repeated KI administrations over 5 days induced urinary elimination of radioiodine over 24 hours (Figure 6C).

Our results indicated that urinary excretion is a major route of elimination of excess iodine. It has been recognized that iodine, a generally scarce element in the diet, is efficiently reabsorbed in the primary urine by the kidneys. Evidence from the literature shows that NIS, expressed at low level in the tubular cells of human kidney, would be involved in these reabsorptions ^68^. NIS immunostaining studies from our group did revealed significant levels of NIS protein expression in the apical membrane of renal tubule cells, suggesting a low NIS expression and, consequently, a saturable absorption of iodide in renal tubule. The effect of the competitive NIS-inhibitor on the uptake kinetics (Figure 5A) indicated the existence of a mechanism of renal reabsorption that has the functional properties of NIS and is therefore most likely mediated by it. The regulation of circulating iodine is thus a priori ensured by a system allowing, on the one hand, a very efficient renal reabsorption of iodine in case of low dietary intake in order to limit its elimination by the urine, and, on the other hand, a rapid urinary elimination of any excess in iodine intake. With an administration of 1 mg/kg per day, the excess iodine was thus mostly eliminated in less than 12 hours in mice. The whole body and the thyroid were a priori sufficiently saturated with iodine to ensure good protection against contamination, without the need to maintain a permanent state of excess circulating iodine. Interestingly, our results also showed that iodide excess saturates the renal reabsorption of circulating iodide and, consequently, increase the urine excretion of circulating radioactive iodide in case of contamination.

### Insights on iodine metabolism in embryos, newborns and lactating rodents

In the present work, using a pin-hole collimator enabled us to perform, for the first time to our knowledge, uptake measurements in mouse embryos.

Low-iodide diets were tested on pregnant mice. Different responses were observed between the organs of the embryo and those of their mother. Under iodine-deprivation, thyroid uptake is markedly reduced in embryos which implies a reduction in thyroid function and in hormone production (Figure 3). Most likely, the mother’s thyroid then produces the hormones for the embryos. This reduction observed in the thyroids of the embryos could also result at least in part from an alteration of placental transfer, but the results obtained on the stomach seem to exclude this hypothesis. Indeed, the accumulation capacity of the radiotracer in the stomach of the embryo did not decrease significantly with the iodine-deprivation. In this context, it is useful to recall that an activation of thyroid function, linked in particular to iodine deficiency, can lead to proliferation of thyrocytes and thus to an increase in the size of the thyroid, and to the formation of a goiter. Some clinical observations indicate an increase in the size of goiters in newborns ^69, 70, 71, 72, 73, 74^ related to low iodine diet but the incidence remains low, especially in cases of moderate deficiency. The fetal thyroid findings reported in this work may contribute to a better understanding of goiter induction during gestation and, in this context, hormone synthesis mainly by mother’s thyroid could explain the low incidence on fetal thyroid.

The incidence of KI treatment was assessed on both pregnant mice and neonates. Our results showed KI administration induced strong decreases of the fetal thyroid uptakes (Figure 8). Considering these strong inhibitions, we then studied the consequences for the neonates of the administration of iodine to the mother before birth (daily gavage) or before and after birth (iodine-enriched diet). A strong reduction of the thyroid uptake capacities of the newborns was observed on the two first days after birth (Figure 9). In parallel, other experiments were performed in rat pups and moderate inhibitory effects were observed on the thyroid after daily KI gavage during maternal gestation (Figure 10). These differences between rats and mice would be interesting to characterize in more detail at the molecular level. In their experiments, Lesbir and collaborators evidenced that repeated KI administration during pregnancy of rats led to long-term irreversible neurotoxicity in the male progeny ^25^. A likely explanation for this stems from the low uptake levels found in neonate thyroids. This lower uptake should lead to a reduction in the synthesis of thyroid hormones by the newborns. At this critical stage of development of the central nervous system, such transient hypothyroidism could be responsible for the observed neurotoxicity.

In lactating animals, a strong decrease in uptake was observed at the mammary gland level following iodine-enriched diet (Figure 8). This result suggests that iodine transport activity in the mammary gland was also regulated by the concentration of circulating iodine through lower NIS expression (Supplementary Figure 3).

In conclusion, our study contributes to a better understanding of the regulation of iodine metabolism at the level of the thyroid and extra-thyroidal organs, including in fetuses and newborns. Extrapolated to the human situation, our data provide guidance on countermeasures against accidental exposure to radioactive iodine and further insights into iodide deprivation as a clinical preparative measure for differentiated thyroid cancer patients.

## Acknowledgments

This work was funded by the French Government under the ‘Investments for the Future’ program (Nuclear Safety and Radioprotection Research (RSNR) action), managed by the French National Research Agency (ANR-11-RSNR-0019)’. We thank Catherine Buchanan for editorial correction of the manuscript.

## Conflicts of Interest

The authors declare no conflict of interest.

---

**Supplementary Figure 1:**
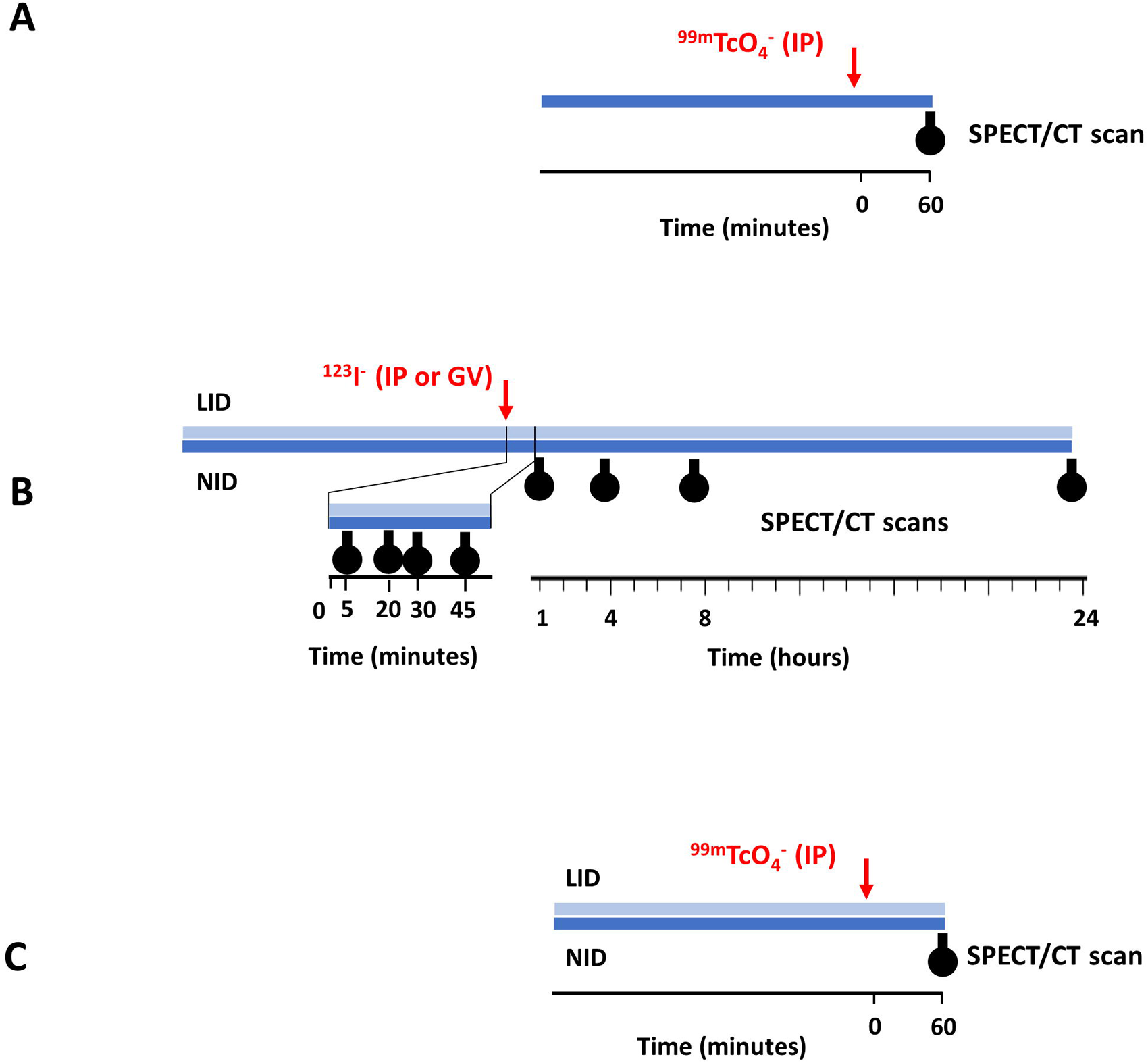
Study protocols for Figures 1 (A), 2 (B) and 3 (C). (A) Mice were fed normal-iodide diet (NID) and radiotracer (^99^mTcO_4-_) was administered intraperitoneally (IP) at time zero. Animals were then anesthetized, positioned on the scanner, and data were acquired one hour after radiotracer injection. (B) Mice were fed either the normal-iodide diet (NID) or low-iodide diet (LID) for 8 days and radiotracer (^123^1-) was administered intraperitoneally (IP) or by gavage at time zero of the kinetic measurements. Animals were then anesthetized, positioned on the scanner, and data were acquired. After one hour acquisition, mice were woken up and put back in their boxes. At indicated times, animals were anesthetized, positioned on the scanner, and data were acquired. (C) Adult control and pregnant mice were fed either the normal­ iodide diet (NID) or low-iodide diet (LID) for 8 days and radiotracer (^99^mTcO_4_-) was administered intraperitoneally (IP) at time zero of the kinetic measurements. Animals were then anesthetized, positioned on the scanner, and data were acquired one hour after radiotracer injection.

**Supplementary Figure 2:**
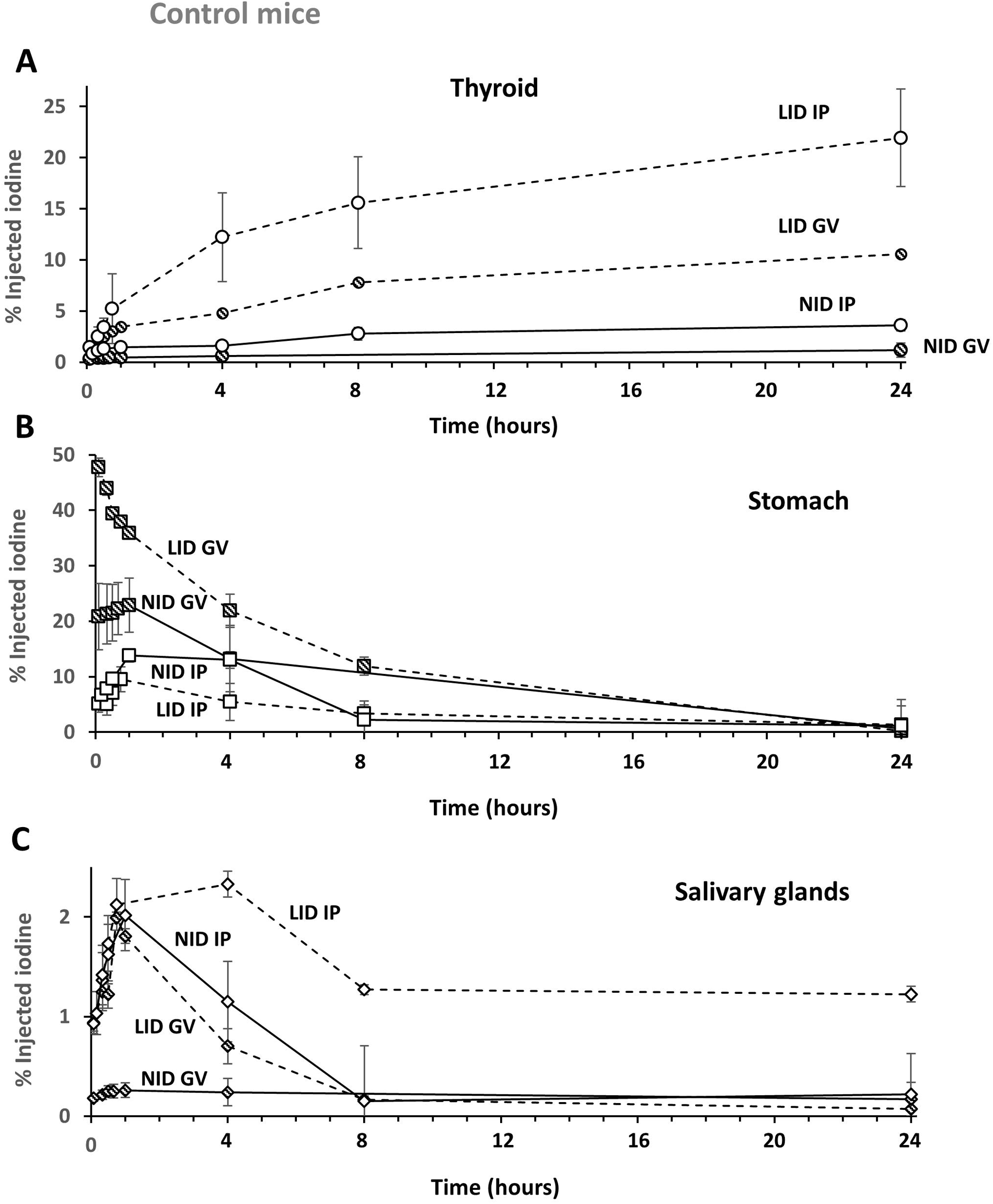
Kinetics of iodine-123 uptake in mouse organs: thyroid (A), stomach (B) and salivary glands (C), as a function of the mode of administration and the iodine diet. The kinetics plotted in dotted line are those obtained with mice under a Low-iodine diet (LID) and in solid line those obtained with mice under a normal-iodine diet (NID). Results obtained by intraperitoneal injection (IP) have white fill marks and those obtained by gavage (GV) have hatched marks. Values have been corrected for radioactive decay and are the average of the percentages of administered doses obtained from at least 5 mice. See study protocol in Supplementary Figure 1B.

**Supplementary Figure 3:**
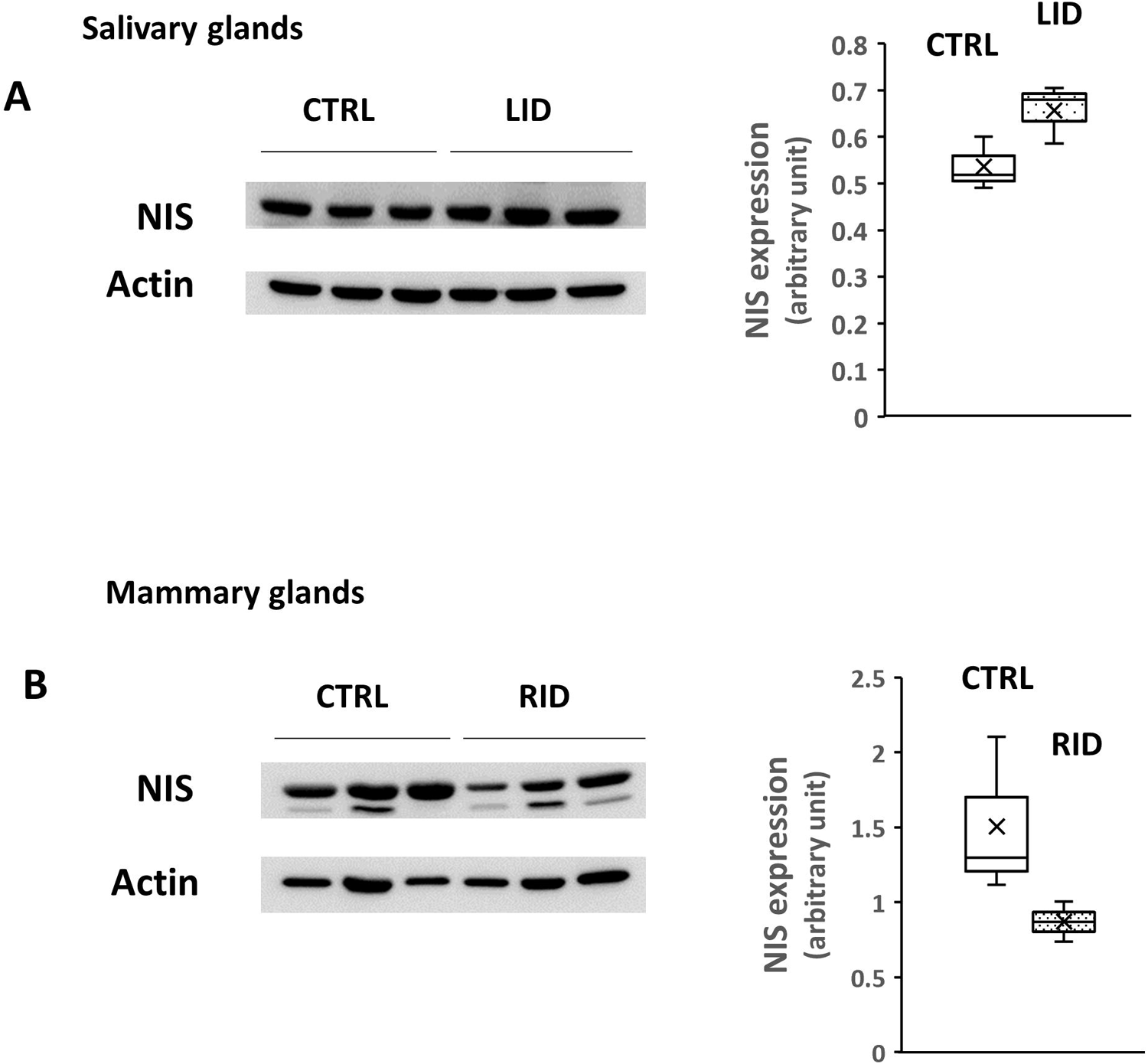
Analysis by Western blot of NIS expression of the stomach (A) and mammary glands (B), as a function of the low iodine diet and the rich iodine diet, respectively. Western blot analyses comparing the effect of different diets on NIS protein expression were performed according to previous publication (Darrouzet *et al*, doi:10.1042/BJ20151086) on total tissue extract using Tris/glycine gels cast in the laboratory. Transfer was performed on the iBlot system from lnvitrogen with PVDF membranes and Tropix I-Block (Life Technologies) as a blocking agent. NIS240 (a monoclonal in-house antibody, the epitope of which is located in the C-terminus region of the NIS protein) was used. The secondary anti-mouse antibody was from Thermo Scientific (reference 31430) and signal detectionwas based on chemiluminescence (GE Healthcare ECL prime). When required, membranes were stripped in 62.5 mM Tris/HCI, pH 6.8, 100 mM 2-mercaptoethanol and 2%SDS for 15 min. Actin protein was probed (monoclonal antibody from Sigma-Aldrich) as a control to verify equivalent protein loading between the variants. Blots were quantified using lmageJ. NIS values have been corrected for actin values and ratios in arbitrary unit are shown (right graphs). Data were obtained from 3 mice for each experimental condition.

**Supplementary Figure 4:**
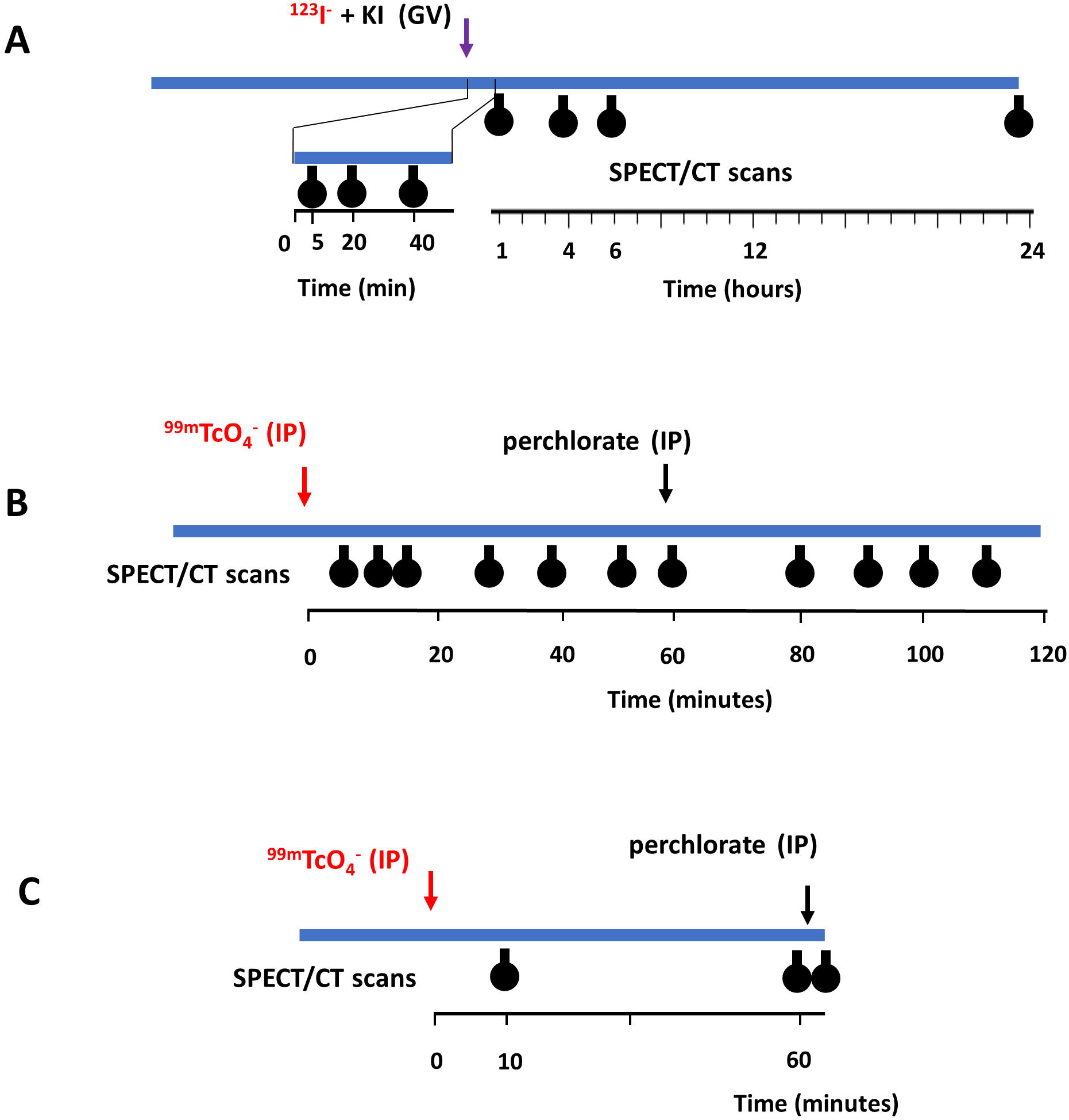
Study protocols for Figures 4 (A), 5 (B, C). (A) Mice were fed the normal-iodide diet (NID). Radiotracer (^123^1-) and Kl (1 mg/kg) was administered by gavage at time zero of the kinetic measurements. Animals were then anesthetized, positioned on the scanner, and data were acquired. After one hour acquisition, mice were woken up and put back in their boxes. At indicated times, animals were anesthetized, positioned on the scanner, and data were acquired. (B) Mice were fed the normal-iodide diet (NID) and radiotracer (^99^mTcO_4_-) was administered intraperitoneally (IP) at time zero of the kinetic measurements. Animals were then anesthetized, positioned on the scanner, and data were acquired. After one hour acquisition, perchlorate was administered intraperitoneally (IP) when indicated, mice were positioned on the scanner, and data were acquired. (C) Mice were fed the normal-iodide diet (NID) and radiotracer (^99^mTcO_4_-) was administered intraperitoneally (IP) at time zero of the kinetic measurements. Animals were then anesthetized, positioned on the scanner, and data were acquired after 10 and 60 minutes. After the one-hour acquisition, perchlorate was administered intraperitoneally (IP) when indicated, mice were positioned on the scanner, and data were immediately acquired.

**Supplementary Figure 5:**
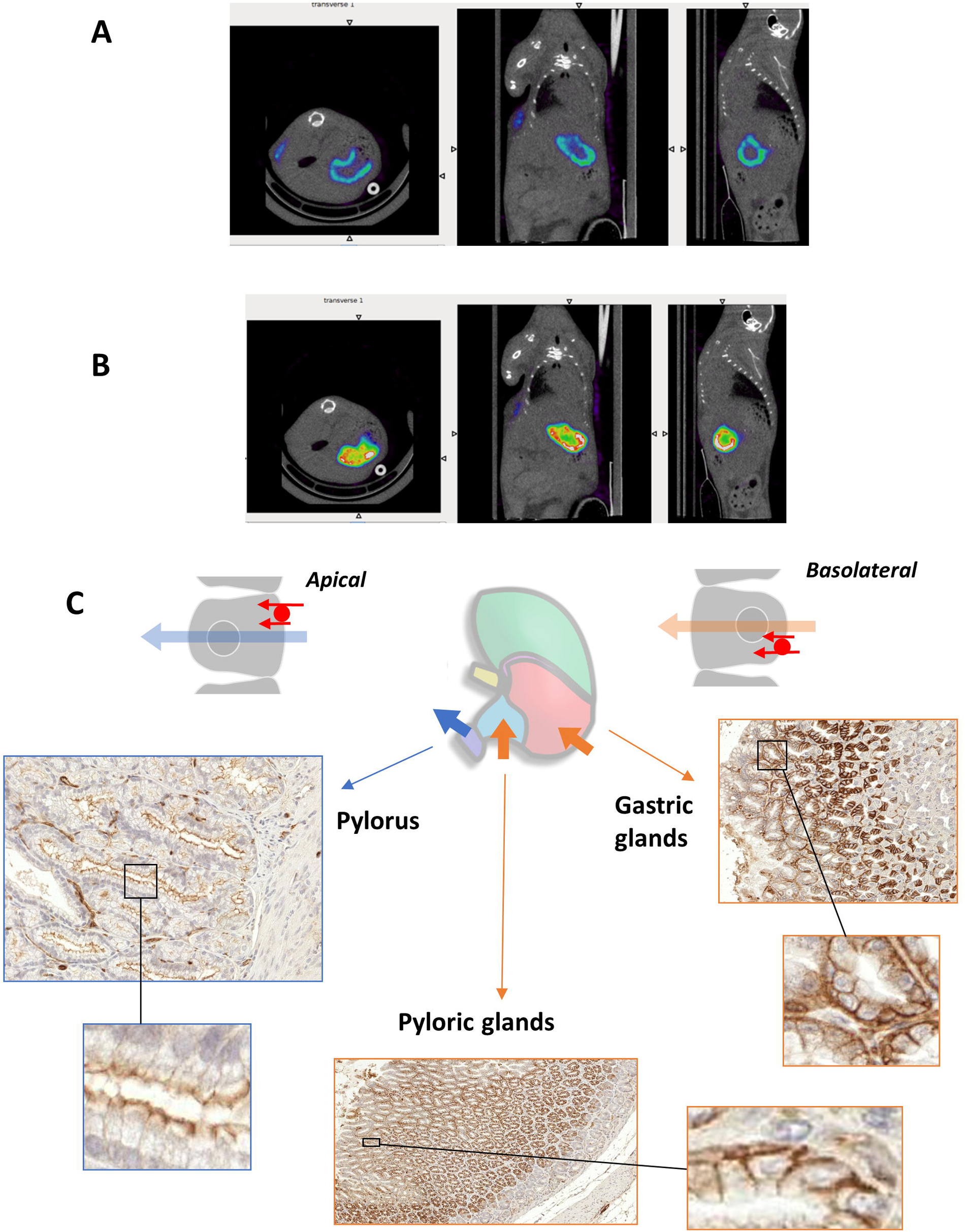
Representative SPECT-CT images of pertechnetate-Tc99m accumulation in the stomach of mice. Animals were imaged 10 minutes (A) and 60 minutes (B) after radiotracer injection. See study protocol in Supplementary Figure 4B. (C) Representative sections of pylorus, pyloric glands and gastric glands of control mice were stained with NIS240, a monoclonal in-house antibody, the epitope of which is located in the (-terminus region of the NIS protein. NIS staining was positive at the apical membrane in pylorus and at the basolateral staining in pyloric and gastric glands.

**Supplementary Figure 6:**
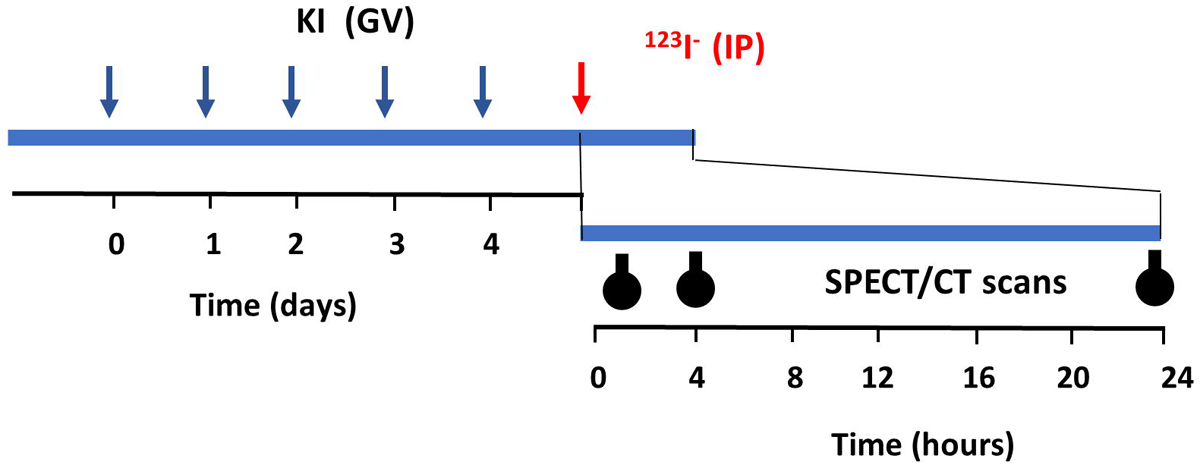
Study protocols for Figures 6. Adult mice and rats were fed the normal-iodide diet (NID). Kl (1 mg/kg) was daily administrated by gavage over 5 days. After 24 hours, radiotracer (^123^1**-)** was administered intraperitoneally (IP) at time zero of the kinetic measurements. Animals were then anesthetized, positioned on the scanner, and data were acquired 1, 4 and 24 hours after radiotracer injection.

**Supplementary Figure 7:**
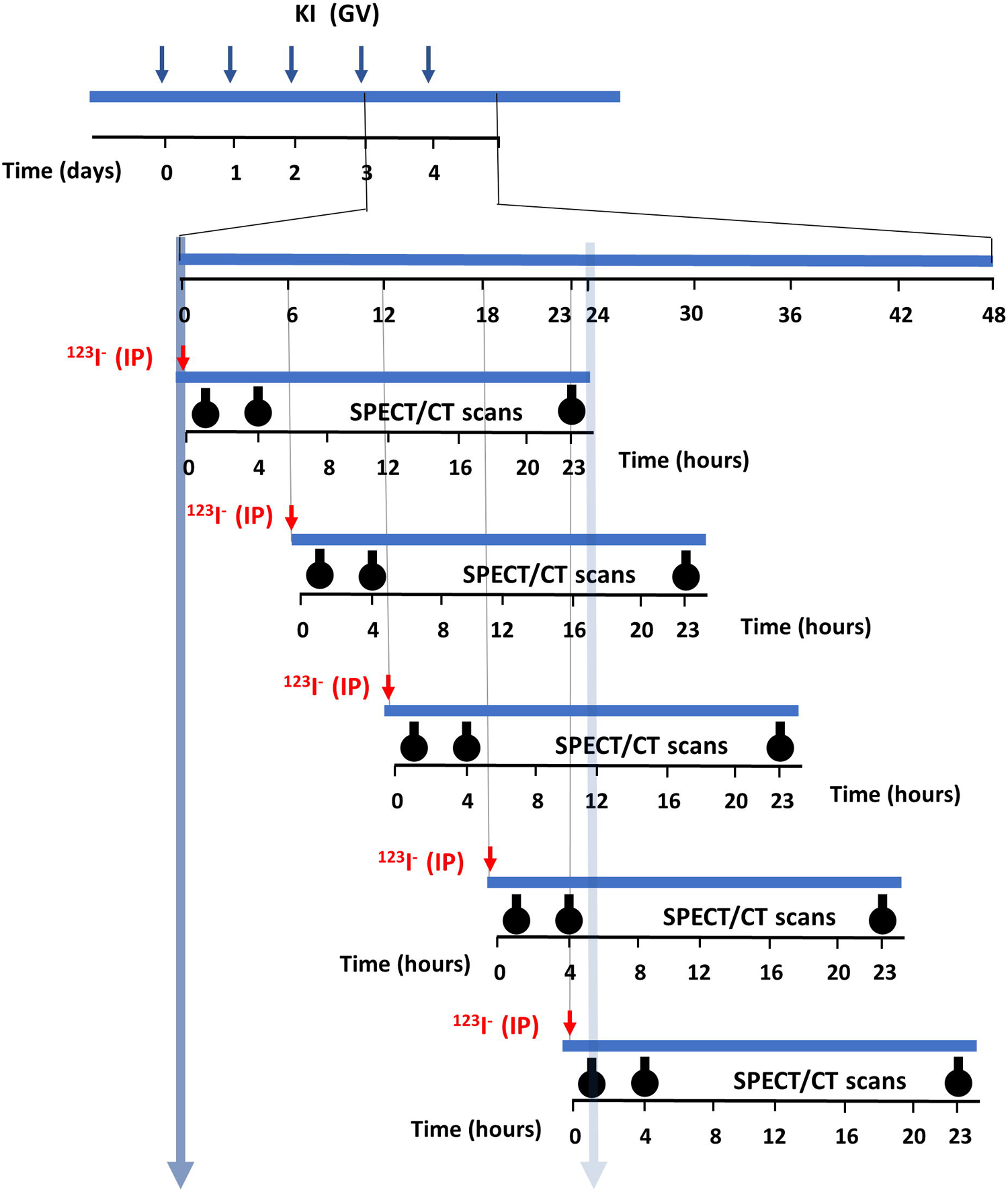
Study protocols for Figures 7. Adult mice and rats were fed the normal-iodide diet (NID). Kl (1 mg/kg) was daily administrated by gavage over 4 days. After indicated hours (O, 12, 18 and 23), radiotracer (1231-) was administered intraperitoneally (IP) at time zero of the kinetic measurements. Animals were then anesthetized, positioned on the scanner, and data were acquired 1, 4 and 23 hours after radiotracer injection.

**Supplementary Figure 8:**
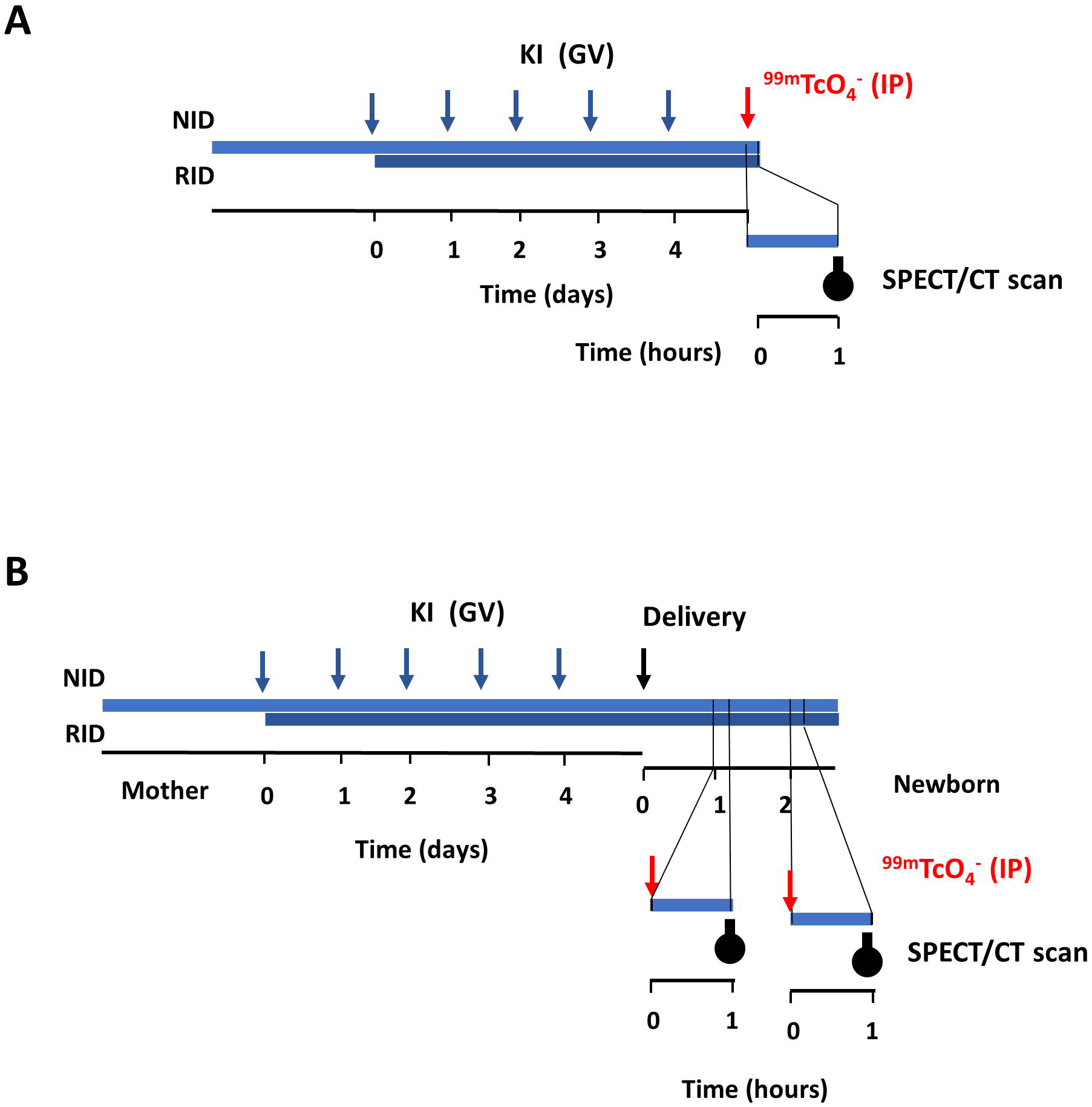
Study protocols for Figures 8, 9 and 10. Adult control, pregnant or lactating mice or rat were fed the Normal-Iodide Diet (NID). Kl was daily administrated by gavage over 5 days (1 mg/kg) (Kl (GV)) or in drinking water (2 mg/L) Rich-Iodide Diet (RID) for 5 days. (A) Radiotracer {^99^mTcO_4_-) was administered intraperitoneally {IP) at time zero of the kinetic measurements. Animals were then anesthetized, positioned on the scanner, and data were acquired one hour after radiotracer injection. (B) Radiotracer {^99^mTcO_4_-) was administered intraperitoneally {IP) to newborns (day one or two after birth) at time zero of the kinetic measurements. Animals were then anesthetized, positioned on the scanner, and data were acquired one hour after radiotracer injection.

## Notes

### Competing Interest Statement

The authors have declared no competing interest.

